# Massively parallel, computationally-guided design of a pro-enzyme

**DOI:** 10.1101/2021.03.25.437042

**Authors:** Brahm J. Yachnin, Laura R. Azouz, Ralph E. White, Conceição A. S. A. Minetti, David P. Remeta, Victor M. Tan, Justin M. Drake, Sagar D. Khare

## Abstract

Confining the activity of a designed protein to a specific microenvironment would have broad-ranging applications, such as enabling cell type-specific therapeutic action by enzymes while avoiding off-target effects. While many natural enzymes are synthesized as inactive zymogens that can be activated by proteolysis, it has been challenging to re-design any chosen enzyme to be similarly stimulus-responsive. Here, we develop a massively parallel computational design, screening, and next-generation sequencing-based approach for pro-enzyme design. As a model system, we employ carboxypeptidase G2 (CPG2), a clinically approved enzyme that has applications in both the treatment of cancer and controlling drug toxicity. Detailed kinetic characterization of the most effective designed variants shows that they are inhibited by approximately 80% compared to the unmodified protein, and their activity is fully restored following incubation with site-specific proteases. Introducing disulfide bonds between the pro-and catalytic domains based on the design models increases the degree of inhibition to 98%, but decreases the degree of restoration of activity by proteolysis. A selected disulfide-containing pro-enzyme exhibits significantly lower activity relative to the fully activated enzyme when evaluated in cell culture. Structural and thermodynamic characterization provides detailed insights into the pro-domain binding and inhibition mechanisms. The described methodology is general and could enable the design of a variety of pro-proteins with precise spatial regulation.

**Significance:** Proteins have shown promise as therapeutics and diagnostics, but their effectiveness is limited by our inability to spatially target their activity. To overcome this limitation, we developed a computationally-guided method to design inactive “pro-enzymes” or “zymogens,” which are activated through cleavage by a protease. Since proteases are differentially expressed in various tissues and disease states, including cancer, these pro-enzymes could be targeted to the desired microenvironment. We tested our method on the therapeutically-relevant protein, carboxypeptidase G2 (CPG2). We designed Pro-CPG2s that are inhibited by 80-98% and are partially to fully re-activatable following protease treatment. The developed methodology, with further refinements, could pave the way for routinely designing protease-activated protein-based therapeutics and diagnostics that act in a spatially controlled manner.

## Introduction

Autoinhibited enzymes, known as pro-enzymes or zymogens, are used ubiquitously for the precise regulation of key biological processes (1). For example, the activities of blood clotting enzymes, digestive proteases, regulatory proteases, apoptotic proteases, and complement system proteins can be triggered by relevant environmental stimuli. Autoinhibiton is typically achieved by extending the active protein with a “pro-segment” or “pro-domain” that inhibits enzymatic activity or prevents adoption of the active conformation until it is cleaved by a tissue-specific activating protease that selectively recognizes a cleavage sequence in the linker connecting the pro-domain to the active protein. To enable tissue specificity for an exogenously administered therapeutic or diagnostic enzyme (2), a general approach for redesigning enzymes into activatable, zymogen variants would be valuable. Surveys of naturally-occurring zymogens have revealed that pro-domains have diverse lengths, folds, and exert their inhibitory effects by steric shielding of the substrate binding sites, or in the case of metalloproteases, by additionally coordinating active site metal ions (1, 3, 4). There have been successful efforts to design pro-enzymes (5–8), but these have relied on unique structural features of the enzyme under consideration. For example, the proximity of N- and C-termini in ribonuclease A allowed for building an inhibiting linker through the active site (5, 6, 8), while barnase allowed for “alternate frame folding”-based design (7) which required proper folding of a chimeric construct containing a duplicated C-terminal segment. Furthermore, like most natural pro-enzymes, these designed zymogens have been limited to enzymes that target macromolecular substrates (5–8), rather than small molecule substrates that are much more difficult to sterically impede and thus represent a significantly greater design challenge. The development of a generalizable, structure-guided approach for zymogen design, particularly one that could be applied to enzymes that target small molecule substrates, would allow spatial control over a broad range of enzymes.

In order to achieve an effective means of localizing the activity of any chosen enzyme to a target microenvironment, we envisioned the addition of an inhibitory pro-domain that would be removable by specific proteases which are markers of the target tissue (Figure 1A). For example, matrix metalloproteases (MMPs) are highly expressed in tumors and would facilitate targeting of enzymatic activity to a tumor site. Tumor-specific MMP activity has been employed previously to achieve site-selective activation of pro-drugs (9, 10) and imaging contrast agents (11). We selected the FDA-approved enzyme, carboxypeptidase G2 (CPG2), as the target for zymogenization. CPG2 is a well-studied and functionally versatile enzyme with a variety of therapeutically beneficial activities (Figure S1). Specifically, CPG2 efficiently inactivates the anti-cancer drug, methotrexate, to reduce systemic toxicity (Figure 1B, Figure S1B) (12–15), for which it has received FDA approval. Moreover, CPG2 activates non-toxic nitrogen mustard prodrugs, such as CMDA (*N*-{4-[(2-chloroethyl)(2-mesyloxyethyl)amino]benzoyl}-L-glutamic acid), to their active, toxic forms in directed enzyme prodrug therapy (DEPT) (Figure S1C) (16, 17). A strategy that exploits the natural properties of tumors to express elevated levels of specific proteases (18, 19) would be broadly applicable for tissue-specific drug delivery. Designed activatable forms of CPG2 which respond to tissue-specific proteases could therefore be useful diagnostic and therapeutic reagents for both delivering and ablating toxicity in a tissue-specific manner (20).

**Figure 1.**
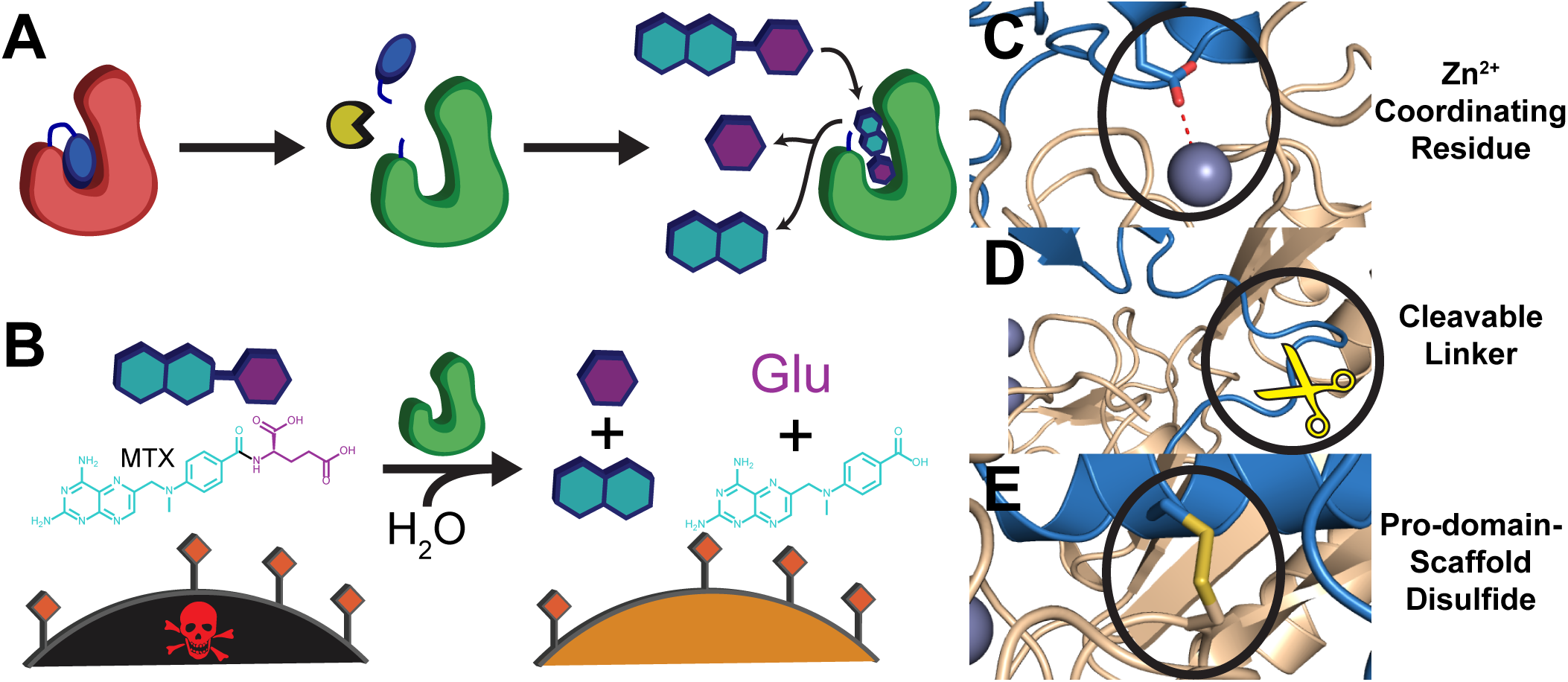
Concept and design strategy for protease-activatable forms of CPG2. (A) Mechanism of activation of Pro-CPG2. A protease (yellow) cleaves the linker that connects the pro-domain (blue) to the inactive enzyme (red), activating it (green). The enzyme can then process small molecules (teal and purple) which can have a biological effect. (B) Activated CPG2 can cleave and detoxify the drug, methotrexate, rescuing the methotrexate-sensitive cells (black to orange). (C) The Pro-CPG2 design strategy ensured incorporation of a zinc (grey sphere)-coordinating glutamate (sticks representation) on the designed pro-domain (blue). The CPG2 scaffold is shown in wheat. (D) The design strategy ensured that the N-terminus of the pro-domain is sufficiently close to the C-terminus of the circular permuted enzyme to incorporate a short linker containing a protease cleavage sequence (yellow scissors). (E) Successful designs were tested for incorporation of a pro-domain-scaffold disulfide bond, designed by Rosetta. All structural figures are Rosetta design models.

Here, we describe a computationally-driven protein design approach to develop protease-activatable zymogens, and employ this strategy to develop pro-enzyme forms of CPG2. Combining next-generation sequencing (NGS) and a computational design approach implemented in the Rosetta software suite, we have developed a massively parallel enzyme design strategy to screen thousands of computationally designed pro-CPG2 variants in parallel. Using this approach, we designed the first artificial pro-enzymes that target a small molecule substrate. We used small-angle X-ray scattering (SAXS) to elucidate the structural basis of inhibition, demonstrating for the first time that key domain movements are critical for the catalytic activity of CPG2. A designed, disulfide-linked pro-CPG2 variant significantly reduces the protection of the PC-3 human prostate cancer cell line from methotrexate-induced cytotoxicity relative to the uninhibited variant. These CPG2 variants are promising starting points for further optimization for anti-cancer therapeutic development and the design methodology may also enable facile zymogenization of other enzymes with therapeutic value.

## Results

### Overview of design strategy

In natural zymogens, pro-domains are fused by a protease-cleavable short linker to one of the protein termini. Inspection of the wild-type CPG2 crystal structure (21) reveals that both termini are >30 Å distant from its active site. We previously designed active circular permutations of CPG2 (CPG2_CP-N89_) with termini proximate to the active site, creating the possibility of attaching a terminal pro-domain that can access and effectively inhibit the active site (22). The CPG2_CP-N89_ variant (in which the polypeptide chain is re-organized such that the N-terminal residue is wild-type residue 89) served as the starting point for our design efforts. As CPG2 is a zinc-containing metalloenzyme, we reasoned that using a zinc-coordinating sidechain in the pro-domain, such as glutamate, to guide pro-domain placement towards the catalytically important zinc ion would effectively block the core active site region when the pro-domain is bound. We thus envisioned appending pro-domains to either terminus such that both a metal-chelating residue and a chosen activating protease cleavage site sequence in the connecting linker are incorporated (Figure 1CD).

### Helical pro-domain design

As an initial proof-of-concept for the auto-inhibition of CPG2, we designed series of pro-enzymes featuring a short helical pro-domain. We used Rosetta to dock a 12- or 17-residue helix into the active site of the CPG2_CP-N89_ circular permutant (22) while allowing substitutions on both the pro-domain and CPG2 sides of the interface. The resulting poses were then subjected to loop closure, fusing the helix to either terminus using a linker containing the TEV protease cleavage sequence. We identified multiple designs in which a Zn^2+^-coordinating residue could be added such that it would interact with the catalytic dinuclear zinc site. When considering the mutations introduced in CPG2, a single substitution, K177A (wild-type residue numbering), was present in many of the designs and appeared to decrease steric restrictions in the formation of pro-domain-enzyme interactions. This substitution only resulted in a ∼20% loss of activity relative to the CPG2_CP-N89_ circular permutant (Figure S2D),which we have previously demonstrated has equivalent activity to the wild-type enzyme (22). Eight designs were selected for activity characterization with either the original TEV cleavage sequence or by substituting the TEV sequence with the MMP-2 cleavage sequence, and two designs (Figure S2B, G and H, Table S1) were experimentally determined to be modestly auto-inhibited (Figure S2). Incubation of pro-enzyme with the relevant protease for 24-48 hours resulted in an enzyme that regained comparable activity to the uninhibited scaffold, and reversion of the K177A mutation resulted in a smaller degree of inhibition when comparing the pro-enzyme and the base variant while only moderately impacting base variant activity. These findings validate the design of a true pro-enzyme that is both inhibited and re-activatable, although the degree of inhibition is modest.

To gain insight into the structural basis for the observed inhibition, we solved the crystal structure of one of the designed pro-enzymes (PDB ID 7M6U, Table S2) and compared it to the structure of wild type CPG2 (PDB ID 1CG2). We could not locate density for the appended helical pro-domain in the structure, a finding that is consistent with the weak inhibition observed for this design. Nonetheless, we were able to determine that this variant maintains the overall tertiary structure of the enzyme (Figure S3, Cα RMSD of 0.26-0.52 Å when comparing all 1CG2 chains to all 7M6U chains), indicating that the circular permutation does not perturb the structure or function of CPG2, as expected from our previous results (22).

### Massively-parallel pro-CPG2 computational design

Building on the modest, proof-of-principle inhibition achieved in the helical pro-CPG2 designs, we pursued a massively-parallel design approach to develop more strongly inhibiting pro-domains. Three characteristics present in the initial designs were used to guide our approach. First, we included an interaction between the catalytic zinc ion and a glutamate residue on the pro-domain as the “hotspot” interaction (Figure 1C). This was achieved by directing pro-domain placement using a library of free glutamate conformations that coordinate the zinc residue; backbone coordinates of these glutamate conformations were used to position the pro-domain. Second, only C-terminal pro-domains were considered, as these proved to be more effective in our initial designs (Figure 1D, Figure S2B). Finally, the K177A mutation, which diminished the enzyme’s activity by ∼20% (Figure S2D) while providing the benefit of decreased steric restrictions at the expected pro-domain binding site, was retained in all designs. A set of 3157 pro-domain scaffolds was generated by combining small domains curated from the PDB and from a large set of previously reported *de novo* designed “miniproteins” (23) (also see Methods). Candidate pro-domains feature a wide range of folds and sequences, including alpha-helical bundles and mixed alpha/beta folds. The pro-domain library can be downloaded at Zenodo (https://doi.org/10.5281/zenodo.5553581).

To minimally perturb the activity of the post-cleavage form of the enzyme, we used a one-sided design approach in which only the pro-domain sequence could be modified while the sequence of the CPG2 “scaffold” region remained fixed. The interaction surface for the pro-domain is not a typical natural protein-protein interaction site, as it includes a significant number of polar and charged moieties such as the dinuclear zinc ion-containing active site. To introduce interactions with these polar groups, we designed explicit hydrogen bond networks between the pro-domain and the enzyme scaffold.

A total of ∼15,000 designs was obtained using a design workflow implemented in Rosetta (Figure 2A; also see Methods). We used a number of criteria (detailed description in Supplementary Results S1, Figure S4, and Table S3) to select ∼7500 designs for testing using a high-throughput assay (as described in the next section). As controls, we included four helical pro-domain designs (one ineffective and three moderately effective autoinhibitors) that allow comparison of the relative activities determined in our high-throughput assay to those of pro-domains with known inhibitory effects.

**Figure 2.**
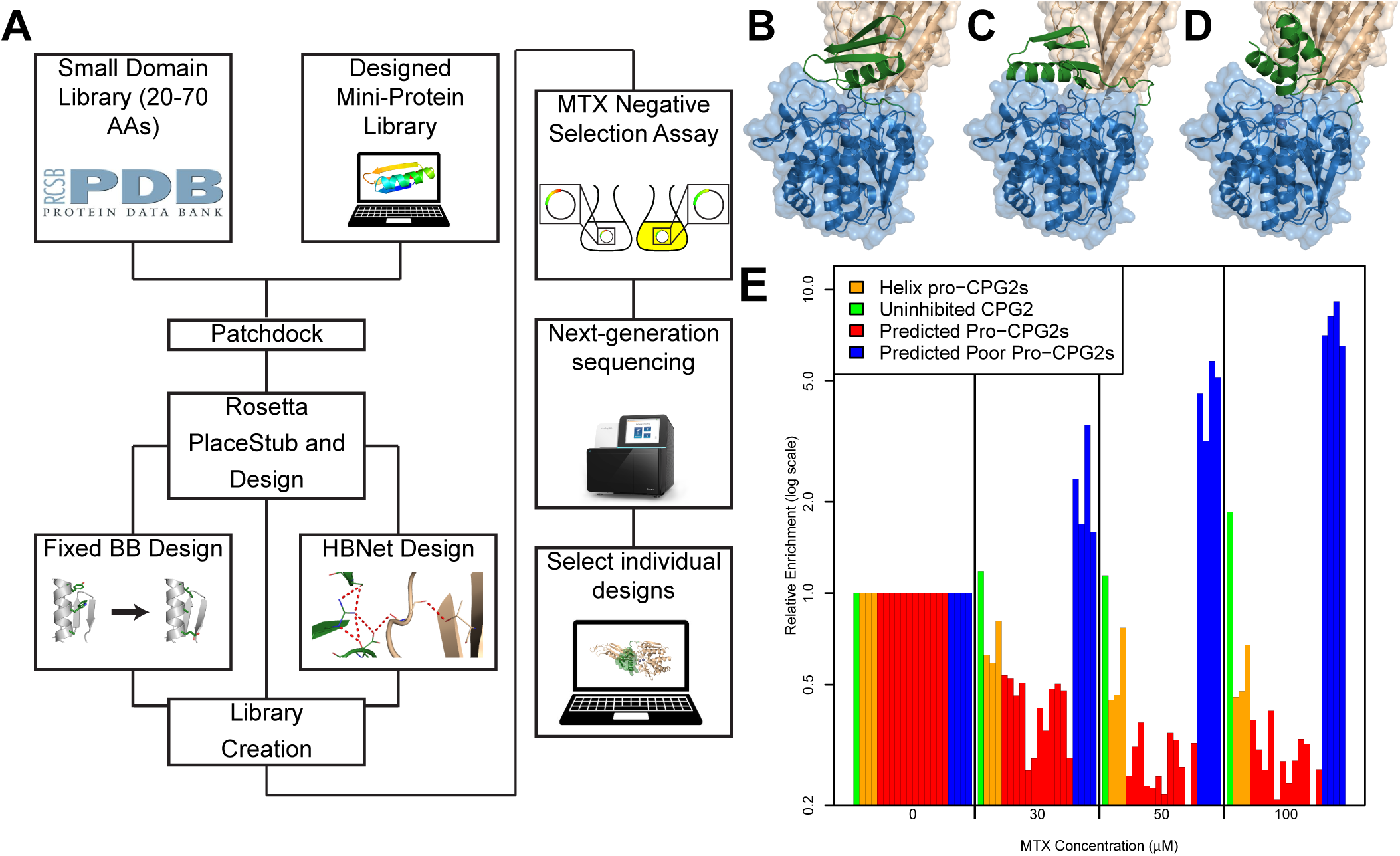
Massively-parallel design of Pro-CPG2. (A) The design strategy is depicted as a flow chart. (BCD) Rosetta models of the three Pro-CPG2 designs selected for detailed study: (B) Pro-CPG2-1, (C) Pro-CPG2-2, (D) Pro-CPG2-3. The pro-domain (green), catalytic domain (blue), and dimerization domain (wheat) are highlighted. (E) Next-generation sequencing results from the massively parallel selection assay. The relative enrichment ratio (log scale) is given at increasing methotrexate concentrations. The helical designs that are known not to inhibit (green) or to mildly inhibit (orange) CPG2_CP-N89_ are shown alongside the twelve designs predicted to be Pro-CPG2s (red) and predicted not to be Pro-CPG2s (blue).

### Massively parallel screening of the pro-CPG2 design pool

We previously reported that periplasmically-expressed CPG2 is able to protect *E. coli* CAG12184 cells from methotrexate toxicity (22, 24–26). We employed this screen to discriminate between inhibiting and non-inhibiting pro-CPG2 designs from the set of 7469 computationally-generated designs. In summary, cells expressing designs with poorly inhibiting pro-domains (i.e. full CPG2 activity) detoxify methotrexate effectively and survive, while cells expressing designs with strongly inhibiting pro-domains (i.e. low CPG2 activity) are unable to detoxify methotrexate and do not survive. We generated a relative ranking of pro-domain efficacy by comparing the fraction of NGS reads attributable to each design in cultures grown at various methotrexate concentrations (30, 50, and 100 μM) and comparing them to cultures grown in the absence of methotrexate. By including both moderately effective and ineffective control pro-domains identified from our original helical pro-CPG2 designs, the NGS results could be analyzed to identify designs that are expected to be at least as strongly inhibited as the helical pro-CPG2s (Figure S2 and Table S4). Between 400,000 and 1.3 million NGS reads were obtained for each individual test culture corresponding to methotrexate concentrations of 0-100 μM, suggesting adequate coverage of the 7500 unique sequences contained in the library.

Upon analysis of the NGS data, reliable enrichment values could be obtained for 1384 out of the tested sequence pool of 7500 sequences, allowing us to identify 107 inhibited designs and 27 “high probability” inhibited designs, the latter defined as designs whose relative enrichment was lower in the 30 µM methotrexate culture as compared to the 0 µM culture, and lower in both the 50 µM and 100 µM methotrexate cultures relative to the 30 µM culture (also see Supplementary Methods). To confirm the ability of this high throughput assay to effectively discriminate between inhibiting and non-inhibiting designs, we selected twelve of these 27 designs, as well as four designs identified to be non-inhibiting (Figure 2E and Figure S5). The twelve designs selected for further study were chosen to maximize sequence diversity and eliminate designs with very similar predicted folds and binding modes. The diversity within the set of inhibiting designs suggests that our selection is not biased towards a particular fold. Each of these sixteen designs was purified as an individual construct alongside the most inhibited helical pro-CPG2 design and the CPG2_CP-N89_-K177A construct. All twelve of the predicted inhibiting pro-CPG2 designs were more strongly inhibited than the helical pro-CPG2 design, while the four predicted non-inhibiting pro-CPG2 designs exhibited activities resembling CPG2_CP-N89_-K177A (Figure S5). These results confirm that our screening system accurately discriminates between inhibiting and non-inhibiting designs, using the helical pro-CPG2 controls as benchmarks. Of the twelve designs, we identified three for further characterization, which we refer to hereafter as Pro-CPG2-1, Pro-CPG2-2, and Pro-CPG2-3 (Table S1). The models of the Pro-CPG2-1 and -2 pro-domains have mixed alpha/beta folds, whereas Pro-CPG2-3 is a helical bundle.

The Michaelis-Menten kinetic parameters of Pro-CPG2-1, -2, and -3 were determined and compared to the fully active CPG2_CP-N89_-K177A (Figure 3ABC and Table S5). Significantly, there is an approximate 50% reduction in the catalytic rate constant (*k*_cat_) for all three designs, indicating a dramatic decrease in the ability of the enzyme to complete its catalytic cycle. In addition, Pro-CPG2-1 and -2 elicited a 2-3-fold increase in the Michaelis constant (K_M_), suggesting that the effective substrate binding affinity is reduced. These three designs exhibit up to an 80% decrease in catalytic efficiency, k_cat_/K_M_.

**Figure 3.**
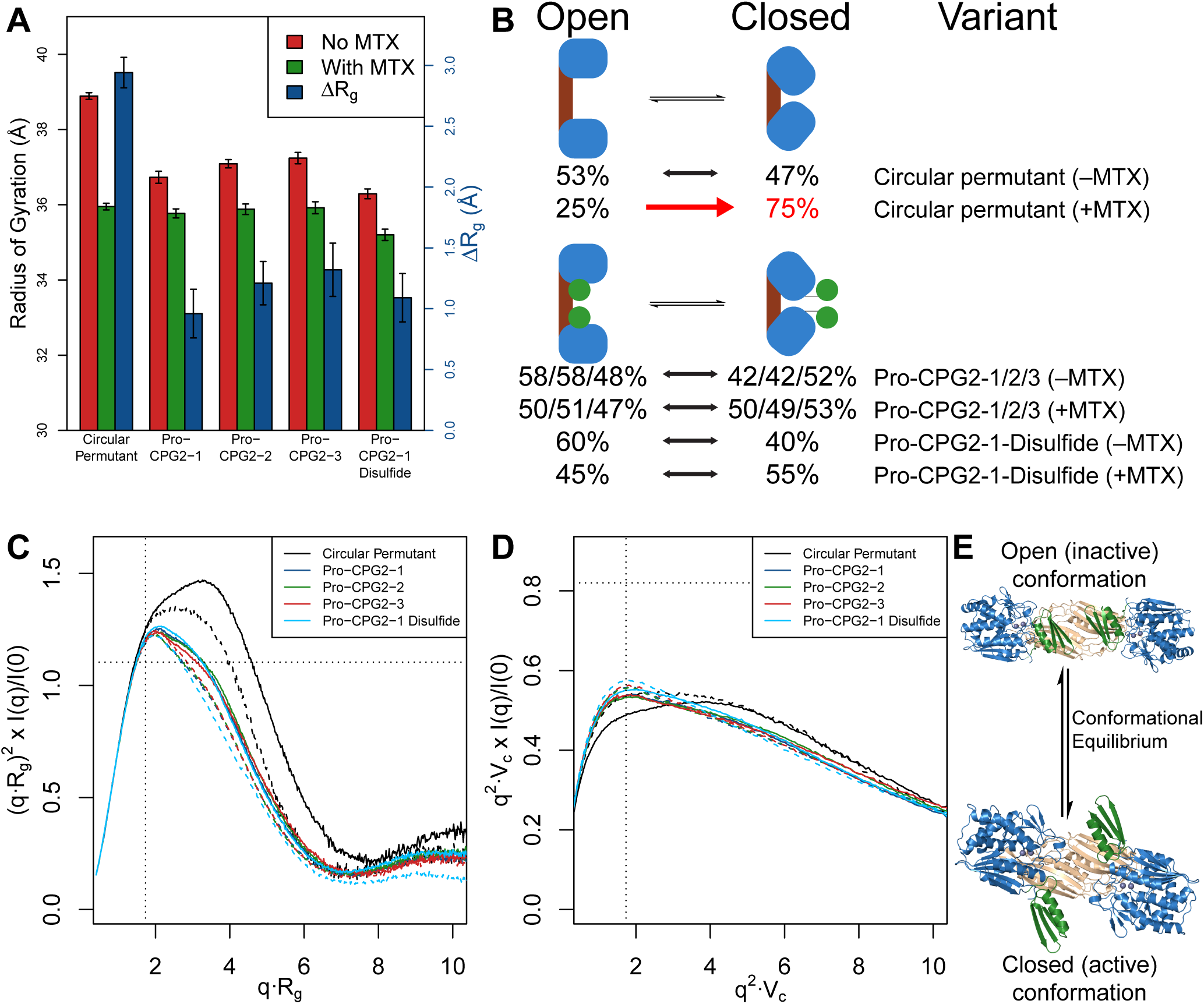
Activity and thermodynamic data for the Pro-CPG2 designs and their disulfide variants. (A-C) The Michaelis-Menten kinetics (A, k_cat_; B, K_M_; C, k_cat_/K_M_) for the three base Pro-CPG2 designs are shown. In yellow, the disulfide variants are grouped with the base design they are derived from. See also Table S5. (D-E) The recovery of molar specific activity following incubation with (D) MMP-2 and (E) TEV protease is shown. Differences in absolute molar specific activity between (D) and (E) are due to different buffer conditions required for MMP-2 and TEV activity. (F) Temperature-dependent thermodynamic binding signatures characterizing the association of CPG2_CP-N89_-K177A and Pro-Domain-1 deduced from ITC measurements. See also Table S7. (G) Temperature-dependent ITC-derived dissociation constant (K_d_) of CPG2_CP-N89_-K177A and Pro-Domain-1. See also Table S7.

### Analysis of inhibited vs. not inhibited designs

Based on the high predictive power of our screening approach, we sought to compare properties of designs identified by our assay to be inhibitors to those identified not to be inhibitors. We performed two-sided *t*-tests comparing the two groups of designs for all computed statistics (selected metrics shown in Table S6). Of note, the number of carbon-carbon contacts between the pro-domain and two regions of interest were found to be significantly greater in the inhibited designs: the base of the dimerization domain beta-sheet (residues 215-219 and 320-324) and a catalytic domain loop that contains the K177A residue previously highlighted in our helical designs and spatially proximal to the circular permutant’s termini (residues 173-182) (Figure S6). These observations suggest that forming contacts with these regions may be important for achieving sufficiently tight binding and inhibition. Upweighting designs with contacts in these regions may yield higher success rates or stronger inhibition in future studies.

### Recovery of activity using specific proteases

In order to function as switches, zymogens must be re-activated by proteolytic processing using a specific protease. To probe the activatability of the pro-CPG2 designs, we generated interdomain linker variants containing the canonical cleavage sequences of two dissimilar proteases: the 3C-type TEV cysteine protease (27) and zinc-dependent matrix metalloproteinase, MMP-2 (28). The activity of all Pro-CPG2 designs is recovered following incubation with the appropriate protease (Figure 3DE). As demonstrated by mass spectroscopy, both residual uncleaved pro-enzyme and retention of the bound but cleaved pro-domain (when free pro-domain is removed through dialysis) may be responsible for the incomplete recovery of CPG2 activity (Figure S7). As two unrelated proteases are able to cleave and re-activate the corresponding CPG2 pro-enzyme, it is likely that a broad range of other site-selective proteases could be used to re-activate pro-CPG2 by including the appropriate cleavage site in the inter-domain linker.

### Characterization of the designed pro-domains

As an initial computational characterization of the designed pro-enzymes, we performed a series of simulations to assess their quality (Supplementary Results S2 and Figure S8). *Ab initio* forward folding successfully “refolds” the designs both as isolated proteins and in the context of CPG2, suggesting that the appended domains are well-folded in the designed conformations.

To further investigate the pro-domains as independent proteins, the pro-domains of Pro-CPG2-1, -2, and -3 were expressed individually and their secondary structure features analyzed using circular dichroism (CD) spectroscopy. All three pro-domains exhibit CD spectra characteristic of well-folded proteins that are stable at physiologically-relevant temperatures. Consistent with the Rosetta models, Pro-CPG2-1 and -2 have characteristic mixed alpha/beta spectra, while the spectrum of Pro-CPG2-3 appears primarily alpha-helical in character. Inspection of thermal melting profiles reveals relatively broad non-cooperative unfolding transitions that do not plateau even when heated to 100ºC, thereby precluding accurate determination of a discrete melting temperature (Figure S9). The high stability of these constructs suggests that they are likely to adopt the same, well-structured fold when fused to the terminus of CPG2_CP-N89_-K177A.

Next, we assessed the binding affinity of Pro-Domain-1 for the CPG2 scaffold “in *trans*.” We conducted isothermal titration calorimetry (ITC) experiments in which Pro-Domain-1 is titrated successively into CPG2_CP-N89_-K177A. Acquisition of a series of binding profiles at varying Pro-Domain (150-500 µM) and enzyme (15-50 µM) concentrations yielded a consistent set of thermodynamic parameters, thereby ensuring that aggregation or precipitation (at higher concentrations) or dissociation (at lower concentrations) do not appreciably impact the binding energetics. Binding isotherms acquired at 5.0ºC reveal that the interaction heats are relatively modest and endothermic. Moreover, analysis of these association profiles indicates that formation of the CPG2_CP-N89_-Pro-Domain-1 complex is characterized by binding affinities on the order of 10^5^ M^-1^ (i.e., *K*_*d*_ ∼ 3-6 µM) over the temperature range studied. Inspection of the thermodynamic binding parameters (Figure 3FG and Table S7) acquired at 5.0ºC reveals that Pro-Domain-enzyme association is entropy-driven, as manifested by the relative contributions of ΔH (1.5 kcal·mol^-1^) and TΔS (8.1 kcal·mol^-1^) to the Gibbs Free Energy (ΔG = -6.6 kcal·mol^-1^).

Significantly, the interaction is accompanied by a slightly negative heat capacity (ΔC_p_ ∼ -0.098 kcal·mol^-1^·deg^-1^) as reflected in the temperature-dependence of binding enthalpies measured at 5 and 15ºC. Based on these results, we anticipate that the enthalpic term approaches zero near 25ºC and is presumably slightly exothermic under selection, assay, and physiological conditions. The observed temperature-dependence is a characteristic hallmark of hydrophobic interactions (29–31) that presumably reflects the desolvation and burial of hydrophobic surfaces, potentially including the methotrexate binding site, upon pro-domain binding.

Examination of the binding interface in our model reveals several non-polar contact regions: 766 Å^2^ of the 1117 Å^2^ buried solvent accessible surface area calculated using Rosetta is hydrophobic. The discrete surface burial, particularly in terms of hydrophobic residues, is generally consistent with a modest heat capacity change estimated for the interaction and may provide further support for our design model. Given that the binding interface includes a metal coordinating glutamate from the pro-domain, it is entirely plausible that water molecules may be dislodged upon association. The zinc-glutamate coordination might therefore contribute additional unfavorable enthalpic contributions as previously reported for protein-metal interactions (32), resulting in an overall net entropy-driven association. An evaluation of the interdependence of the CPG2 pro-domain binding sites is included in the Supplementary Results (S3).

### An open-closed equilibrium associated with CPG2 activity revealed by small-angle X-ray scattering

In the absence of a crystal structure where the binding mode of our designed pro-domains could be identified, we sought to use small-angle X-ray scattering (SAXS) experiments to understand the structural states occupied by our designs as compared to the uninhibited enzyme (Figure 4). We hypothesized that, similar to other members of the M20 family of metallodipeptidases, CPG2 exists in two distinct conformations: an “open” conformation responsible for substrate binding and a “closed” conformation optimized for catalytic turnover (Figure 4E). The former conformation is observed in the crystal structure of CPG2 reported here (Figure S3), as well as a previously published structure of CPG2 (21). The latter conformation is observed in many closely-related homologues such as mouse carnosinase 2 (mCN2) (33), Sapep (34), PepV (35), peptidase T (36), aminoacylase-1 (37), and *N*-succinyl-L,L-diaminopimelic acid desuccinylase (DapE) (38). Anticipating that the addition of excess product (which is converted entirely from the substrate by the enzyme prior to data collection) would push the conformational equilibrium to the “closed” conformation, we acquired SAXS data in the presence and absence of hydrolyzed methotrexate to monitor changes in the accessibility of these two states. A model of the “closed” state was generated by individually superimposing the CPG2 catalytic and dimerization domains onto the structure of mCN2, and using generalized kinematic loop closure (39) to re-build the interdomain linkers.

**Figure 4.**
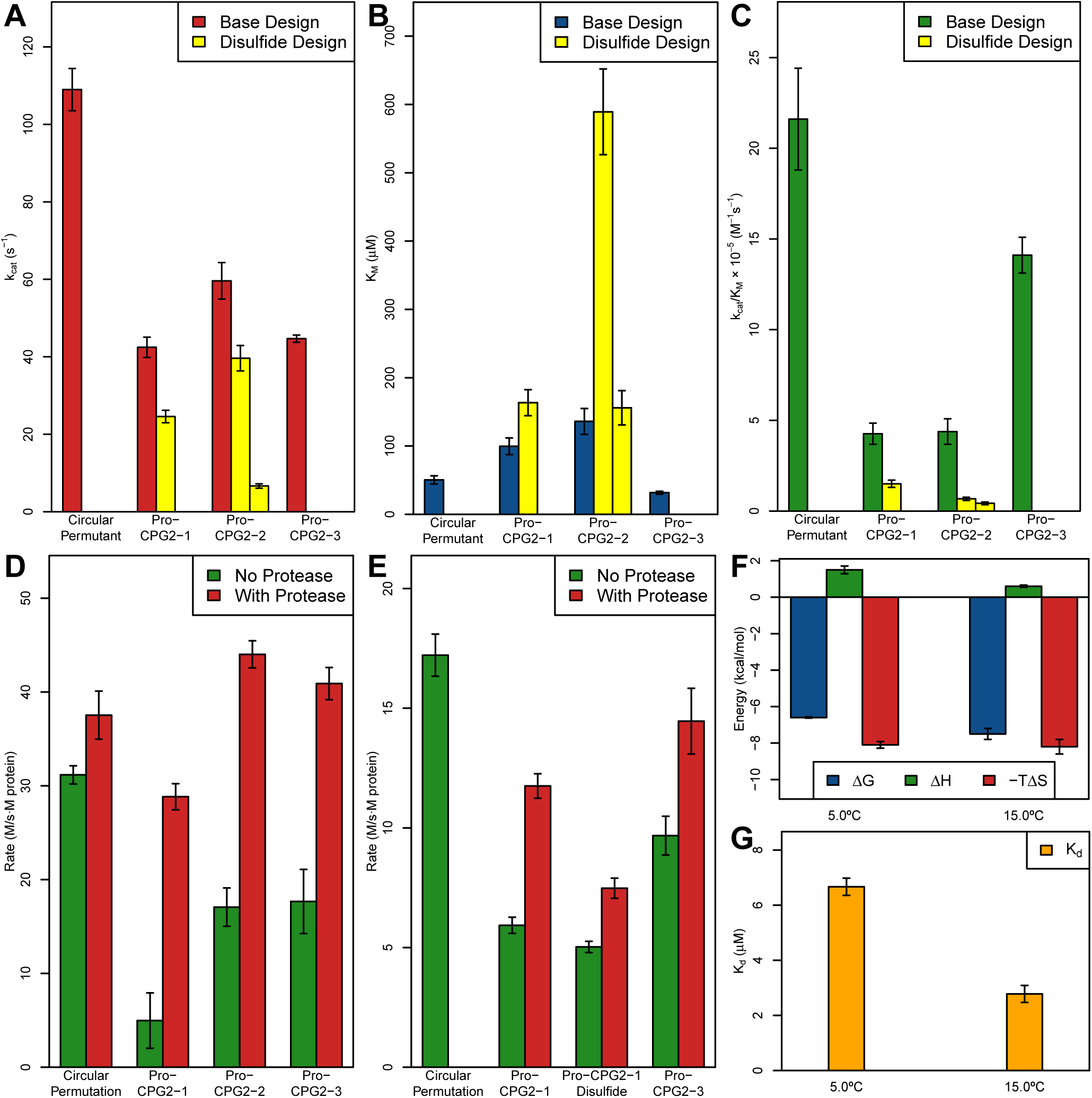
Small angle X-ray scattering (SAXS) analysis for CPG2_CP-N89_-K177A and the Pro-CPG2 designs. (A) The radius of gyration of the enzymes in the absence (red) and presence (green) of hydrolyzed methotrexate (left axis) and difference in radius of gyration (blue, right axis) are shown. (B) Results from fitting the two-state model (open and closed) to the SAXS data using FoXS are shown, with “-MTX” and “+MTX” indicating the absence or presence of hydrolyzed methotrexate. The quality of the fits is shown in Figure S12. (C-D) The (C) radius of gyration-based and (D) volume of correlation-based dimensionless Kratky plots are shown. Samples with solid lines have no hydrolyzed methotrexate present, while samples with dashed lines have hydrolyzed methotrexate present. The horizontal and vertical dotted lines indicate cross at the peak position of a compact, spherical particle. In the R_g_-based plot, if the peak is shifted up and to the right, this indicates that the particle is more elongated, and if the curve becomes a hyperbolic plateau, this is indicative of a disordered particle. In the V_c_-based plot, a downwards shift indicates an increase in surface-to-volume ratio relative to that of a sphere. (E) Rosetta models of the conformational equilibrium required for CPG2 activity, highlighting the role of the pro-domain (green).

Analysis of the SAXS data (Figure 4A, Table S8) for CPG2_CP-N89_-K177A in the absence of the products resulted in a particle with a radius of gyration (R_g_) of 38.9 Å. While neither the “open” crystal structure nor the “closed” model produced a reasonable fit of the data alone (χ^2^ > 25), we observed a good fit (χ^2^ = 2.2) for a two-state model using FoXS (40). The two states (“open” and “closed”) each contribute roughly 50% of the total scattering (Figure 4B). In contrast, when hydrolyzed methotrexate is added to the sample, one obtains a smaller R_g_ of 36.0 Å which is consistent with the more compact “closed” model (Figure 4A). The “closed” model produced a reasonable fit to the data using CRYSOL (41) (χ^2^ = 4.9), though a moderately better fit is obtained with the two-state FoXS model (χ^2^ = 2.3), with 75% and 25% contributions from the “closed” and “open” conformations, respectively (Figure 4B). These data support our hypothesis and are consistent with the mechanisms of homologous enzymes (33–38), namely that addition of the product triggers a shift in the conformational equilibrium towards the “closed” conformation of CPG2.

We conducted a similar analysis on Pro-CPG2-1, -2, and -3 (Figure 4 and Table S8), using Rosetta-generated models of open and closed states of these proteins. In all three cases, the R_g_s of samples containing substrate are reduced when compared to samples in the absence of product; however, the magnitudes of these changes are smaller than those for CPG2_CP-N89_-K177A, primarily owing to a smaller observed R_g_ for samples without product (Figure 4A).

When two-state models of the Pro-CPG2s are used to fit the scattering data, the conformational equilibrium is approximately 50%/50% open/closed, as observed for CPG2_CP-N89_-K177A, ranging between 48-59% in the “open” conformation. In contrast, the conformational equilibrium does not shift with the addition of product, remaining at roughly 50%/50% levels for the pro-enzymes (Figure 4B). It should be noted that while the precise percentages of populations of the two states are highly dependent on the quality of the models used for fitting the data and should not be interpreted in *absolute* terms, the lack of a shift towards the “closed” conformation in *relative* terms is consistent, regardless of the models used. Thus, SAXS-based characterization of all Pro-CPG2 variants is consistent with a scenario in which the “open” state is preferentially stabilized by the designed inhibitory domain.

To further corroborate our findings and investigate the impact of bias introduced by the quality of both the CPG2_CP-N89_-K177A and pro-enzyme models, we utilized model-free analyses of the SAXS data, including dimensionless Kratky (42, 43) and Porod (42) exponent analyses (Supplementary Results S4 and Figure 4). These analyses confirm that in the absence of product, CPG2_CP-N89_-K177A behaves as a well-folded but flexible particle. While addition of the product renders this particle less flexible, the pro-enzymes’ flexibility is even further reduced, with the concentration of mass occurring closer to the particle’s center of mass. These results are therefore consistent with our initial findings that the pro-enzymes exhibit reduced conformational flexibility as compared to the circular permutant, with a greater population adopting the inactive, “open” state.

### Effects of disulfide stapling on Pro-CPG2

To validate our structural models of Pro-CPG2 and to establish an “upper bound” of enzyme inhibition using our current designs, we generated disulfide-stapled variants of Pro-CPG2-1 and -2 with the goal of restricting the pro-domain to its designed “open”-state conformation. For each pro-enzyme, we used a Rosetta-based approach to identify locations where a disulfide bond spanning the CPG2-pro-domain interface is feasible (Figure 1E, Figure S10). We identified one disulfide variant of Pro-CPG2-1 and two variants of Pro-CPG2-2 that exhibited substantially reduced activity (Figure 3ABC) relative to the CPG2 cysteine variant (Table S1). Notably, when the other three CPG2 cysteine variants (without the pro-domain) generated by Rosetta (of a total of six) were tested, they resulted in enzymes that were catalytically inactive. In other words, all disulfide pro-enzyme variants designed by Rosetta that were compatible with the structure and function of CPG2 resulted in reduced activity. SAXS experiments on the Pro-CPG2-1 Disulfide variant reveal a further moderate decrease in flexibility as deduced from the dimensionless Kratky plots (Supplementary Results S4 and Figure 4CD). We conclude that a disulfide bridge is effectively created between the pro-domain and scaffold, thereby validating our structural models. Moreover, “pinning down” the pro-domain using a secondary covalent linkage assists in strengthening the effectiveness of the pro-domain, presumably by decreasing its conformational freedom, and establishes a reasonable maximum inhibition that could be obtained following optimization of the Pro-CPG2-1 design.

### Pro-CPG2-1-Disulfide has reduced ability to detoxify methotrexate in cancer cell culture

As we plan to use the designs described here as a starting point for the development of therapeutics or diagnostics, we sought to ascertain the level of inhibition in a more complex biological context prior to initiating the extensive optimization required for specific applications. To that end, we treated an aggressive prostate cancer cell line, PC3, with Pro-CPG2-1-Disulfide and CPG2_CP-N89_-K177A. The PC3 cell line was derived from a bone metastasis of a grade IV prostate cancer patient and is commonly used to assess treatment responses, including chemotherapies, making it ideal for evaluating the capability of designed pro-enzymes to degrade the chemotherapeutic methotrexate in a cell survival assay (44). As methotrexate is toxic to PC3 cells, CPG2 activity in the cell culture medium is expected to increase cell survival in the presence of methotrexate. Designed pro-CPG2 enzymes have significantly lower methotrexate hydrolysis activity, and are, therefore, expected to lead to decreased cell survival compared to unmodified CPG2. PC3 cells were incubated with Pro-CPG2-1-Disulfide or CPG2_CP-N89_-K177A and then subsequently treated with 50 nM methotrexate over six days. Cells treated with the Pro-CPG2-1-Disulfide variant indeed displayed significantly lower dose-dependent cell viability at enzyme concentrations of 15-62 pM when compared to CPG2_CP-N89_-K177A (Figure 5). These data indicate that the designed disulfide-containing pro-enzyme remains effectively autoinhibited in a cell culture-based assay. When a similar experiment was performed with Pro-CPG2-1, which is less strongly inhibited but could allow proteolytic removal of the pro-domain, a smaller, qualitative reduction in PC3 cell viability was also observed as compared to CPG2_CP-N89_-K177A (Figure S11). Nonetheless, the increased cell survival of the disulfide variant suggest that further optimization of the Pro-CPG2-1 design is possible and may yield effective switches in selected therapeutic contexts.

**Figure 5.**
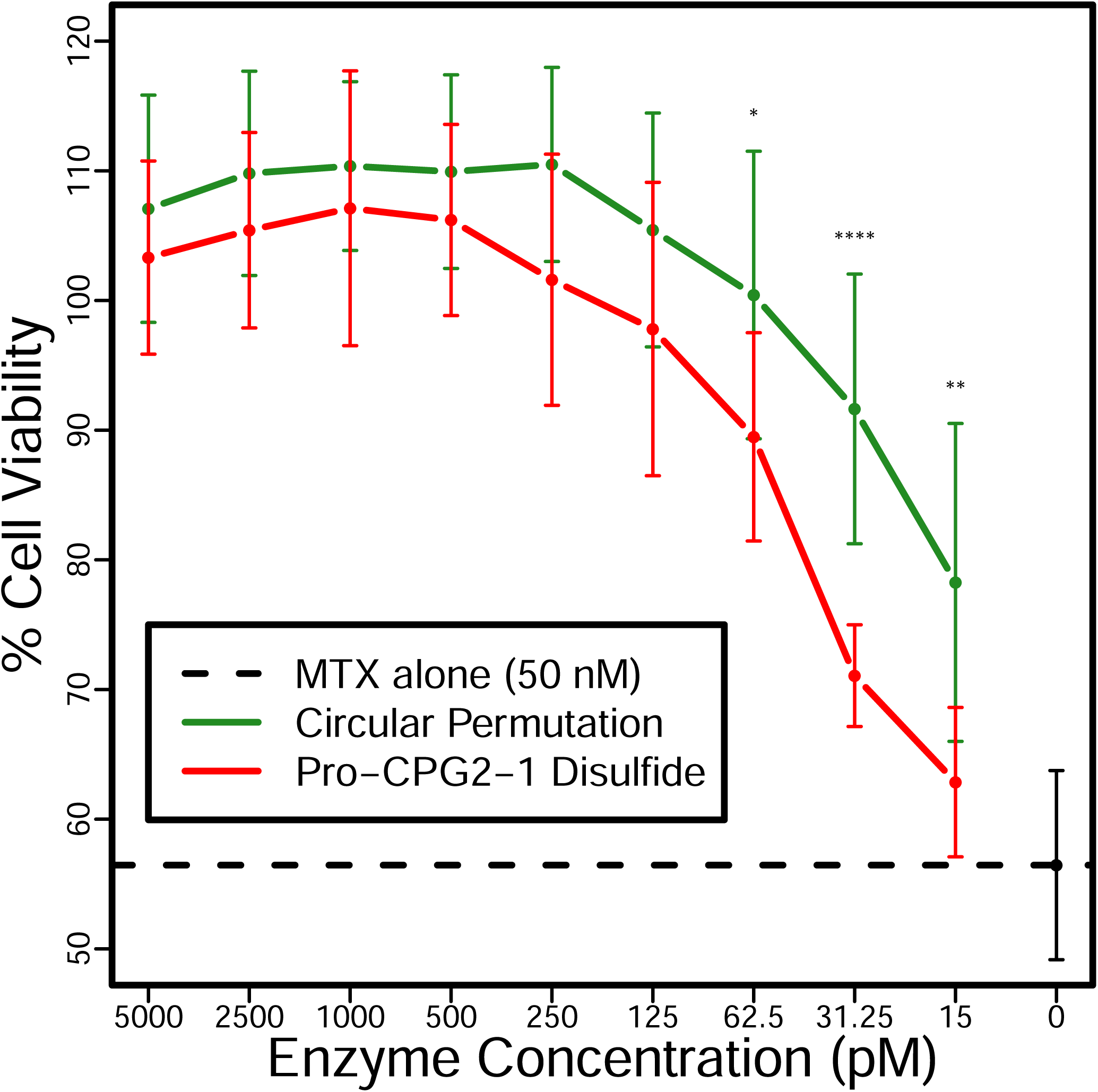
Pro-CPG2-1-Disulfide has reduced ability to cleave and detoxify methotrexate (MTX) compared to CPG2_CP-N89_-K177A. PC3 cells were treated with MTX at 50 nM, and then treated with either CPG2_CP-N89_-K177A or Pro-CPG2-1-Disulfide. Enzyme concentrations between 250-5,000 pM showed 100% viability. Enzyme concentrations between 15-62 pM showed a significant difference (unpaired t-test, p<0.05) of viability in PC3 cells between the two enzyme treatment groups, with cells treated with Pro-CPG2-1-Disulfide exhibiting a lower survival rate than cells treated with active CPG2. The dashed line indicates the cell viability in the presence of MTX alone (no enzyme treatment). Error bars indicate the standard deviation. *= p< 0.05, **= p< 0.01, ****= p < 0.0001. Two-factor analysis of variance p < 10^-8^. See also Figure S11.

## Discussion

We have developed a massively parallel enzyme design and screening approach to engineer a pro-enzyme form of CPG2, an FDA-approved drug and one of the best-studied DEPT enzymes. This approach has allowed us to achieve a reversible decrease in activity of up to 80% relative to the CPG2_CP-N89_-K177A variant without further design optimization. Inhibition can be reversed by the activity of two structurally and functionally distinct proteases, including MMP-2 that is overexpressed by many cancer cell lines. Introduction of a disulfide bond between the pro-domain and the enzyme, which allows us to simultaneously validate our structural model and establish an upper bound for inhibition for our design, resulted in 98% inhibition compared to the base CPG2_CP-N89_-K177A variant although the reactivation of disulfide-containing protein is less efficient.

The design challenge for developing a pro-CPG2 enzyme is substantial. Unlike natural and previously engineered zymogens which act upon macromolecular substrates, CPG2 substrates are relatively small molecules, offering less opportunity to directly sterically impede substrate binding by design. The pro-domain binding site is not known to naturally form protein-protein interactions whose binding modes can serve as templates for computational design. Moreover, considering its proximity to the active site, any substitutions within the catalytic or dimerization domains could decrease the activity of the post-cleavage form of the designed pro-enzyme.

Therefore, we employed a one-sided design strategy in which only the pro-domain is designed while the rest of the protein sequence is held fixed, further increasing the challenge as compared to design strategies in which both partners in a protein-protein interface are allowed to be optimized simultaneously.

Computational approaches using *de novo* designed mini-proteins have shown great promise in the development of protein binders against a variety of therapeutically relevant targets (23, 45), but these methods have not been used thus far for the design of pro-enzymes. The ability to synthesize genes corresponding to tens of thousands of designed proteins as oligonucleotide pools coupled with a high-throughput screening assay and next-generation sequencing allow testing a broad swathe of sequence and structure space for the desired functionality, allowing for successful design campaigns in spite of low success rates (<10%) (45) in the computational design step. We used these developments to design mini-protein-based pro-domains. Although the individual candidate mini-protein scaffolds are predicted to be well folded in isolation, success in pro-enzyme design requires that they fold properly and remain soluble and stable when fused to the CPG2 circular permutant. To serve as an effective pro-domain, they should feature sufficiently robust inhibitory interactions while still allowing dissociation upon protease cleavage. Furthermore, as is common for enzymes, multiple states (open and closed) are likely involved in the catalytic cycle, but high resolution structures are available for only one (open) state, highlighting the additional challenges unique to pro-enzyme design compared to protein-protein interface design. Given these considerations, we reasoned that a massively parallel design and screening approach with a diversity of pro-domain structures would maximize opportunities to identify strongly inhibiting pro-domains that can also be effectively removed by proteolysis.

It is noteworthy that the pro-domain structures and sequences that we identified are highly varied, suggesting that the design approach developed here is able to identify multiple, distinct solutions to the problem, rather than converging on a single solution, as is often the case with “winner-takes-all” directed evolution strategies.

Compared to natural zymogens in which activity level differences between inhibited and activated states are ∼100-fold, the best (non-disulfide-containing) designed CPG2 zymogens show ∼5-fold increases upon activation, indicating that these diverse solutions identified by computation require further improvements, possibly using directed evolution, in future efforts aimed at improving the dynamic range of the designed pro-enzymes for therapeutic applications. One avenue for improvement would be to allow further substitutions in the catalytic domain that may improve pro-domain binding. A comparative analysis of computational designs that are and are not effective inhibitors also provides insights for further improvement (Figure S6, Table S6). Contacts between the pro-domain and specific regions in the catalytic and dimerization domains appeared to be significantly greater in designs identified as inhibitors in the massively parallel screening assay, suggesting that increasing the focus on these regions may yield stronger inhibitors during design optimization.

There are at least two putative mechanisms for the inhibition of activity observed in Pro-CPG2 designs. In one case, if the pro-domain blocks the active site or binds nearby, it may impede binding of the substrate, either directly or through subtle distortions of the active site.

Alternatively, as CPG2 is expected to undergo a “closing” conformational change during catalysis, the pro-domain may act as a “door stop,” preventing the enzyme from closing after substrate binding, thereby impeding catalysis. Because we designed the pro-domains to be in close proximity to the catalytic zinc ions while also having them located at the “hinge region” of the expected closure site, our design strategy could impact either or both of these mechanisms of inhibition. Our structural and kinetic characterization provides insight into the mechanisms of inhibition. If the pro-domain is acting to block the active site, rather than directly interfering with the catalytic machinery of the enzyme, one might expect that this would manifest as a decrease in affinity with the substrate, as evidenced by an increase in K_M_. As the catalytic residues and conformational changes are unimpeded in this model, the k_cat_ should be minimally affected. Conversely, if the substrate is free to bind the enzyme, but the enzyme’s ability to pass through its catalytic cycle is impeded, the k_cat_ is expected to be reduced, while the K_M_ should be unaffected. In two of the three most inhibited designs, we observe an increase in K_M_ and a decrease in k_cat_, suggesting that both mechanisms are at play. The third design is characterized solely by a decrease in k_cat_, suggesting that the primary mechanism of inhibition occurs *via* disruption of catalytic turnover.

The SAXS data provide further insight into the molecular mechanism of inhibition. We present the first direct evidence that CPG2 undergoes open-closed conformational changes, although similar observations have been reported for homologous enzymes (33–38). Our data highlight the conformational selection underlying CPG2 activity: CPG2_CP-N89_-K177A, upon addition of product, shifts towards the more compact, “closed” conformation. In the Pro-CPG2 designs, this conformational shift does not seem to occur, suggesting that this change is impeded by the presence of the pro-domain. This trend is further validated by analysis of the SAXS flexibility data; while various SAXS-derived metrics suggest that CPG2_CP-N89_-K177A is more compact and less flexible in the presence of hydrolyzed methotrexate than in its absence, the Pro-CPG2 samples display minimal corresponding variation. Furthermore, these data are generally consistent with a more compact Pro-CPG2 structure.

Analysis of the calorimetric profiles characterizing in-*trans* binding of the pro-domain of Pro-CPG2-1 revealed dissociation constants in the low micromolar range. In a therapeutic context, this affinity is sufficient to inhibit the enzyme, while likely being weak enough to be readily released from the binding site following proteolytic processing. In order to evaluate whether pro-enzymes with similar inhibitory characteristics have potential utility for *in vivo* or clinical applications, we examined whether the pro-enzyme form of CPG2 has diminished ability to protect PC-3 cancer cells from methotrexate toxicity. Our results demonstrate that the decreased activity of the Pro-CPG2-1-Disulfide variant results in a statistically significant reduction in the enzyme’s ability to detoxify the drug over a multi-day experiment in tumor cells, while the “base” Pro-CPG2-1 variant shows a smaller, qualitative difference in cell survival. This activity differential suggests that further investigation of these pro-enzymes on additional cell lines and animal models as well as in clinically valuable contexts is warranted. For example, the use of a tumor activated pro-enzyme able to activate a chemotherapeutic pro-drug in antibody DEPT (ADEPT) would further mitigate the serious side-effects associated with conventional chemotherapy beyond what is already achieved using ADEPT. ADEPT enzymes such as CPG2 are directed to the tumor using a targeting molecule, such as a tumor-directed antibody, where they can activate the drug, thereby localizing the cytotoxic effects to the tumor site. Clearance times for the circulating enzyme vary widely from patient to patient (46, 47), requiring the development of a variety of complex methods for assaying serum enzyme levels (46) and ultimately precluding their clinical use to date. While we previously designed a light-activated, pro-drug-activating enzyme (48), challenges associated with the low tissue penetrance of light may limit the applicability of this approach in the clinic as compared to protease-activated enzymes. As the Pro-CPG2 enzymes that we have developed exhibit reduced activity compared to their activated form in tumor cell culture, selection of an appropriate protease that is upregulated in the tumor microenvironment could facilitate synergistic improvement in the pharmacological safety profile of ADEPT therapy. The ability to easily switch the activating protease offers a distinct advantage, as the approach can be readily adapted to tumors with varied protease expression profiles.

## Conclusion

The computational design of zymogen forms for any chosen enzyme is a challenging problem, particularly for enzymes acting on small molecule substrates. This study presents a massively parallel computational design and screening approach to develop diverse pro-enzymes of CPG2, a clinically useful enzyme acting on small molecule substrates, whose inhibition is readily reversed through proteolytic cleavage of the enzyme-pro-domain linker. A key aspect of our computational method is the use of a structurally diverse set of pro-domains to create novel protein-protein interfaces, and our screening approach exploits advances in synthetic DNA technology and NGS to build and screen a library of thousands of pro-domains without requiring specialized assays or equipment. The characterization of our best pro-enzyme design reveals that two mechanisms of inhibition are at play: binding of the substrate to the enzyme is impeded, and domain movements required for catalytic activity are also disrupted. Finally, we demonstrate that the activity differential between the disulfide-linked, inhibited form and activated form is sufficient to significantly impact cell survival under methotrexate treatment, while the design without the disulfide link showed a smaller impact on cell survival. This design and screening strategy has potential implications for the development of a broad range of enzyme inhibitors and pro-enzymes, potentially yielding novel therapeutics and diagnostic tools.

## Materials and Methods

Complete experimental methods are described in the Supplementary Information.

### Hotspot placement and pro-domain design

Rosetta’s “PlaceStub” mover (49) was employed to place small domains interacting with the catalytic zinc ion. An interaction between one of the active site zincs and a glutamate introduced in the pro-domain was defined as the “hotspot.” A library of 466 inverse rotamers of glutamate interacting with the active site zinc was built using the “TryRotamers” mover. Hotspot-directed pro-domain placement was performed using script 3 (Table S9). This protocol produced 4444 designs, which were further filtered to 1668 designs after analyzing the scores and filter values.

### Increasing pro-domain sequence diversity

To generate a larger set of designs, further rounds of fixed backbone design were applied to the set of 4444 designs produced from the Hotspot protocol. Either the standard fixed-backbone design protocol or a design protocol (HBNet) aimed at obtaining hydrogen bond networks at the interface (50, 51) was used as described in scripts 4 and 5 (Table S9). These protocols produced 8757 (fixed backbone) and 4993 (HBNet) additional designs, for a total of 15,418 designs (Table S4). As described in the Supporting Information, 7500 designs and controls were ordered and screened using a high-throughput cell survival assay.

### Disulfide variant design

Disulfide-bridge variants were designed starting from the base pro-domain models using the Disulfidize mover (39) (script 8, Table S9) and evaluated for quality manually.

### Determination of CPG2 molar specific activity

CPG2 was first diluted to 100 μg/mL in 50 mM Tris, 100 mM NaCl, pH 7.4, and then to 1-10 μg/mL in the same buffer, after which 10 μL was mixed with 90 μL of 50 mM Tris, 50 mM NaCl, 0.1 mM ZnSO_4_, pH 7.4 containing 100 μM of methotrexate pre-heated to 37ºC (final protein concentration of 100-1000 ng/mL). The absorbance at 320 nm was measured over time in a 96-well plate (Greiner Bio-One UV-Star Half Area 96-well Microplate) using a Tecan Infinite M200 Pro. Absorbance units were converted to units of molar concentration of methotrexate, and initial rates (molar concentration of methotrexate consumed per second) were determined using linear regression. Activity is reported as the initial rate divided by the molar concentration of CPG2. In cases where relative activity is reported, the molar specific activity is divided by the molar specific activity of the reference enzyme and converted to a percentage.

### Michaelis-Menten kinetics

CPG2 was first diluted to 100 μg/mL in 50 mM Tris, 100 mM NaCl, pH 7.4, and then to 50 ng/mL, or 500 ng/mL for the disulfide variants, in the same buffer. 10 μL of this sample was mixed with 90 μL of 50 mM Tris, 50 mM NaCl, 0.1 mM ZnSO_4_, pH 7.4 containing between 1-300 μM methotrexate, pre-heated to 37ºC. The final protein concentration was approximately 1 nM (or 10 nM for the disulfide variants). The absorbance at 320 nm was measured over time as described above. Absorbance values were converted to methotrexate concentrations, and initial rates were determined using linear regression. Michaelis-Menten parameters were determined using non-linear regression.

### CPG2 Cell Viability Assay

PC-3 cells were seeded at 1,400 cells/100 μL in 96 well plates in RPMI media (Sigma-Aldrich) with 10% FBS, 1% Pen/Strep, and 1% GlutaGro (Corning). After an overnight incubation at 37°C and 5% CO_2_, media in the wells was replaced with fresh RPMI media with 5% charcoal-stripped (CSS)-FBS, 1% Pen/Strep, and 1% GlutaGro (Corning). Cells were then grown for 3 days. We then added one of the following three drug groups: 50 nM methotrexate alone, 50 nM methotrexate + CPG2_CP-N89_-K177A (100-5,000 pM), or 50 nM methotrexate + Pro-CPG2-1-Disulfide (100-5,000 pM). Treatment lasted for 6 days with replenishment of the media, drug, and enzymes after 3 days. Cell viability was measured using WST-1 at 1:10 dilution with CSS-FBS media at absorbance of 450 nm (Tecan 1100 Plate Reader).

## Supporting information

Supp lnfo

## Acknowledgements

This work was supported by a grant from the NIH (R01GM132565 to SDK). BJY was supported by a Canadian Institutes of Health Research Post-Doctoral Fellowship and a Mistletoe Research Fellowship. LRA was a recipient of a RosettaCommons Research Experience for Undergraduates internship program. VMT was funded by the National Institute of General Medical Sciences (NIGMS) Grant T32 GM135141. We would like to thank Joseph D. Bauman and Eddy Arnold (Rutgers University Center for Advanced Biotechnology and Medicine) for providing support and resources for the crystallography experiments, and Vikas Nanda (Rutgers University Center for Advanced Biotechnology and Medicine) for providing access to an Aviv Model 420SF CD Spectrometer. We would like to acknowledge Dr. Dibyendu Kumar, Dr. Min Tu, and the Waksman Genomics Core Facility for performing the NGS analysis. SAXS data were collected at the Advanced Light Source (ALS), SIBYLS beamline on behalf of US DOE-BER, through the Integrated Diffraction Analysis Technologies (IDAT) program. Additional support comes from the NIGMS project ALS-ENABLE (P30 GM124169) and a High-End Instrumentation Grant S10OD018483. Mass spectroscopy data were collected by Amenah Soherwardy and Haiyan Zheng (Rutgers University Biological Mass Spectrometry Facility). The authors acknowledge the Office of Advanced Research Computing (OARC) at Rutgers, The State University of New Jersey for providing access to the Amarel cluster and associated research computing resources that have contributed to the results reported here.

## Author contributions

BJY and SDK designed the project. BJY, CASAM, DPR, JMD, and SDK designed the experiments. BJY performed all experiments and analysis of data unless otherwise indicated. LRA performed the computational design of the first-generation helical pro-enzymes. REW and VMT performed the PC-3 cell survival assays and analyzed the data. CASAM and DPR performed the ITC experiments and analyzed the data. All authors contributed to the writing and/or editing of the manuscript.

