## Supplementary material for "Massively parallel, computationally-guided design of a pro-enzyme": Supp lnfo

PNAS

### Supplementary Information for
#### Massively parallel, computationally-guided design of a pro-enzyme

Brahm J. Yachnin[1,2], Laura R. Azouz[1,2], Ralph E. White III[3], Conceição A. S. A. Minetti[1], David P. Remeta[1], Victor M. Tan[2,6], Justin M. Drake[3,4,5], Sagar D. Khare[1,2,*]

[1]Department of Chemistry & Chemical Biology and the [2]Institute of Quantitative Biomedicine, Rutgers, The State University of New Jersey
[3]Department of Pharmacology, [4]Urology, and [5]Member, Masonic Cancer Center, University of Minnesota
[6]Department of Pharmacology, Robert Wood Johnson Medical School, Piscataway, New Jersey, USA.

*To whom correspondence should be addressed:
Sagar D. Khare
Center for Integrative Proteomics Research, Room 208M
Rutgers, The State University of New Jersey
174 Frelinghuysen Road
Piscataway, NJ 08854-8076
(848) 445-5143


**This PDF file includes:**

Supplementary Table of Contents
Supplementary text
Figures S1 to S12
Tables S1 to S10
SI References

Figure S7. Mass spectrometry analysis of Pro-CPG2-1 before and after proteolytic cleavage. 20
Figure S8. Computational assessment of the model quality of (A) Pro-CPG2-1, (B) Pro-CPG2-2, and (C) Pro-CPG2-3. 21
Figure S9. Circular dichroism spectra for the three Pro-Domains and their melting curves. 22
Figure S10. Rosetta models of the three disulfide designs. 23
Figure S11. Pro-CPG2-1-Disulfide has reduced ability to cleave and detoxify methotrexate (MTX) compared to $CPG2_{CP-N89}$-K177A. 24
Figure S12. SAXS data and the two-state model fits for circularly permutated CPG2 and the pro-domains. 25
**SUPPLEMENTARY TABLES** 26
Table S1. Sequences of experimentally-characterized Pro-domains. 26
Table S2. Crystallographic data for the $CPG2_{CP-N89}$-K177A-MMP-Helical Pro-domain Crystal Structure (PDB ID 7M6U). 27
Table S3. Computational metrics used to filter pro-CPG2 design candidates. 28
Table S4. Summary of next-generation sequencing statistics. 28
Table S5. Michaelis-Menten parameters for the circular permutation of CPG2, the three Pro-CPG2 variants, and the disulfide variants. 29
Table S6. Differences in computationally computed metrics between predicted inhibited and predicted non-inhibited designs. 29
Table S7. Summary of the thermodynamic binding profiles for the $CPG2_{CP-N89}$-K177A: Pro-Domain-1 complex. 29
Table S8. SAXS data parameters. 30
Table S9. rosetta_scripts referred to in the text. 31
Table S10. Primers used in molecular biology. 32
**SUPPLEMENTARY REFERENCES** 34

#### SUPPLEMENTARY MATERIALS AND METHODS

**Helical pro-domain design.** To assess the feasibility of auto-inhibiting CPG2 using a pro-domain, an idealized helical backbone structure was used. Either two or three repeats of the helical "EAAAK" sequence were fused and placed in roughly the correct orientation to form either an N- or C-terminal connection with the $CPG2_{CP\text{-}N89}$ circular permutant (1). Docking was performed using script 1 (Table S9). As a means to "filter" the resulting placements of the helix, Rosetta Match (2, 3) was used to build a zinc-coordinating residue (Cys, Asp, Glu, or His) between the helix and the zinc coordination site ordinarily occupied by substrate (4). For designs in which co-ordination was deemed possible, steric clashes between the pro-domain and catalytic domain were relieved by further sequence and structure optimization (script 2, Table S9). Favorably-scoring designs were identified, and eight "consensus designs" were selected to be tested experimentally.

**Generation of the $CPG2_{CP\text{-}N89}$-K177A and Helical $CPG2_{CP\text{-}N89}$-K177A-Protease-Pro-domain constructs.** Starting from the previously described $CPG2_{CP\text{-}N89}$ constructs (N- and C-terminally His-tagged) (1), the K177A mutation was introduced using standard Kunkel mutagenesis (5) or QuikChange mutagenesis (Agilent Technologies Inc.) protocols (Table S10, primer 1). To the C-terminal His-tag construct, a C-terminal TEV protease cleavage sequence was inserted upstream of the His-tag using primer extension PCR (Table S10, primers 2-3) followed by QuikChange mutagenesis. This construct was linearized at the C-terminus (Table S10, primers 4-5), and helix design sequences generated by primer extension PCR were inserted using Gibson assembly (6) (Table S10, primers 6-13). The N-terminal, thrombin cleavable His-tag variants were generated by amplifying the entire coding region (Table S10, primers 14-15), excluding the His-tag, and inserting it using Gibson assembly (6) into pET15b digested using NdeI and XhoI (New England Biolabs Inc.). The TEV cleavage site was substituted for an MMP cleavage site using primer extension PCR (Table S10, primers 16-17) followed by QuikChange mutagenesis.

**Small domain library preparation.** Following the success with the simple helical designs, a library of larger pro-domain candidates was prepared from small protein domains. The PDB was screened for high-quality crystal structures of 20-70 residue protein chains (resolution < 2.5 Å). Only monomers were considered, and a maximum of 70% sequence identity between two proteins in the library was allowed, yielding 258 small domains.

In addition, a large set of 40 residue *de novo* designed, well-folded "mini-proteins" (7) were included. Designs that had a stability score (7) of greater than 1.0 were included in our library, yielding an additional 2899 small domains with four different folds.

The structures of all 3157 domains were energy-optimized using the REF2015 scorefunction in Rosetta.

**Initial placement using PatchDock.** To globally dock the library of small domains with CPG2, PatchDock (8, 9) was used. A Rosetta-optimized model of the circularly permuted $CPG2_{CP\text{-}N89}$ dimer was set as the receptor. The K177A mutation was introduced in the starting model, as this mutation had only marginal effects on catalytic activity while improving the degree of inhibition in the helical pro-domain designs. The binding site included residues 114-115, 175-178, 200-201, 324, 360-362, and 384-385 (wild-type CPG2 numbering). A distance constraint of 25 Å between the C-terminus of $CPG2_{CP\text{-}N89}$-K177A and the N-terminus of the pro-domain was used. The top 50 models for each pro-domain were used in subsequent stages.

**Pro-CPG2 library design.** We intended to screen 7500 sequences, including 31 controls, requiring us to reduce our 15,418 designs down to 7469 based on computational scoring metrics. First, designs were excluded based on filter cutoffs as described in Table S5. Designs in which the pro-domain clashed with its symmetric partner and duplicate sequences were also removed, leaving 7511 designs (1256 designs from the original set, 2856 from the fixed backbone design set, and 3399 from the HBNet design set). The poses with the worst Rosetta total_scores from each design set were dropped, until a total of 7469 designs remained. These sequences, plus 31 controls known to inhibit or not inhibit CPG2 from the original helical design strategy, were codon-optimized using Biopython (10). PCR adapters were added (5' adapter: 5'-CTTTATTTTCAAGGGGGGTCT; 3' adapter: TAAAAGCTTAATTAGCTGAG-3'), and these sequences were ordered as an Oligonucleotide Library (Agilent Technologies Inc.) (see https://github.com/BYachnin/pro_cpg2_design_scripts/blob/main/generate_sequences.py).

**Pro-CPG2 library construction.** Starting with $CPG2_{CP\text{-}N89}$ with an N-terminal pelB leader sequence in the pQE-80L expression vector, the K177A variant was prepared as described above to generate pelB-$CPG2_{CP\text{-}N89}$-K177A. This construct was linearized at the C-terminus after incorporating a C-terminal TEV protease cleavage sequence (Table S10, primers 5 and 18). The parental DNA was digested using DpnI.

The Oligonucleotide Library was amplified using the standard Herculase II Fusion PCR protocol (Agilent Technologies Inc.), using 15 PCR cycles (Table S10, primers 19-20). The PCR product was purified. In a 20 µL total volume, 100 ng each of the linearized vector and library of inserts were combined with NEBuilder HiFi DNA Assembly Master Mix and incubated at 50ºC for 1 hour. NEB 10-beta Electrocompetent *E. coli* (50 µL) were transformed with 2 µL of the resulting reaction product. After allowing the cells to recover for 1 hour at 37ºC in 1 mL of SOC, the transformants were expanded into 10 mL of LB media with 100 µg/mL ampicillin and grown overnight at 37ºC. The plasmid DNA was purified and will be referred to as the “Full Plasmid Library.”

**Massively parallel library screening.** The Full Plasmid Library DNA was transformed into electrocompetent CAG12184 cells (11), obtained from the Yale Coli Genetic Stock Center (CGSC #7437). The transformants were expanded into 10 mL of LB media with

100 µg/mL of ampicillin and 15 µg/mL tetracyclin and grown overnight at 37ºC. The cells were resuspended in 50 mL of fresh media and grown again overnight at 37ºC.

Multiple 50 mL cultures of M9 minimal media containing 0.4% glucose, 0.2% casamino acids (made from pure amino acids), 100 µg/mL ampicillin, 15 µg/mL tetracyclin, and 10 mM isopropyl β-D-1-thiogalactopyranoside were inoculated with 1 mL of the library culture. Triplicate library cultures containing 0 µM, 30 µM, 50 µM, or 100 µM methotrexate were prepared alongside control cultures inoculated with CAG12184 cells harboring pQE-80L plasmids with pelB-CPG2$_{CP\text{-}N89}$-K177A, pelB-mCN2, or the helical pro-domain controls. These cultures were grown at 30ºC for 14 hours. Plasmid library DNA from each culture was purified and will be referred to as the “Selected Plasmid Libraries.”

The 12 Selected Plasmid Libraries and the Full Plasmid Library were amplified using 10-12 cycles of the standard Q5 polymerase PCR protocol (Table S10, primers 19-20). These PCR products, as well as the Oligonucleotide Library, were prepared for Illumina NGS. Samples were amplified with an additional 8 PCR cycles to incorporate DNA barcodes using the Illumina Nextera XT kit. A Bioanalyzer 2100 was used to confirm library size and quantity. Samples were mixed in an equimolar ratio and sequenced on the MiSeq instrument using a 300 bp cycle that yields ~10 million reads. After demultiplexing and removing adapter sequences, forward and reverse reads with a minimum 15 bp overlap with 10% error were joined using Flash (12).

Sequences were aligned using Magic-BLAST (13) (FASTQ sequences as queries and the Oligonucleotide Library sequences as our subject database). Alignments flagged as being unmapped or a secondary alignment were discarded, as were alignments with insertions, deletions, skipped regions, hard clipping, padding, or mismatches. Sequences with soft clipping outside of the beginning and end of each sequence or longer than 70 bases each were discarded. The counts of the remaining matches to each of the designs in the Oligonucleotide Library were converted to percentages of the total number of matched sequences present in each NGS sample. Across replicates, the average, standard deviation, and percent error of the percentages for each design was calculated (see https://github.com/BYachnin/pro_cpg2_design_scripts/blob/main/deep_seq_magicblast.R). Computational metrics and NGS-derived data were deposited to ProtaBank (https://www.protabank.org/study_analysis/BhKCzXma/).

To exclude unreliable sequencing data, we filtered designs according to four criteria: the average percentage of the total number of sequences for that design needed to be at least $10^{-3}$% in both the (1) Full Plasmid Library and (2) Selected Plasmid Library with a methotrexate concentration of 0. (3) The percent error in these libraries needed to be less than 50% for the Selected Plasmid Library with a methotrexate concentration of 0, and (4) less than 40% for the Full Plasmid Library. This reduced the number of designs under consideration from 7469 to 1384 (Table S4).

Strongly inhibiting pro-CPG2 designs should decrease as a fraction of the number of sequences present in a culture as the methotrexate concentration increases. To identify such designs, an "enrichment ratio" was calculated by dividing the percentages for a design in the Selected Plasmid Libraries (with varying methotrexate concentrations) by the percentage of that same design in the Selected Plasmid Library with a methotrexate concentration of 0. We identified any design that had an enrichment ratio lower than that of all the helical pro-CPG2 controls as a "Predicted Pro-CPG2" design candidate. The 27 designs in which the enrichment ratio at 50 μM and 100 μM methotrexate were both less than the enrichment ratio at 30 μM methotrexate were identified as "High Probability Pro-CPG2" design candidates. These designed were inspected individually, and twelve were selected for expression as individual designs (Figure 2E, Figure S5, and Table S4). In addition, four designs that were expected to be poorly inhibited were also expressed (Figure 2E and Figure S5). Non-inhibiting designs were identified using the inverse comparison as for the inhibiting designs: the enrichment ratio needed to be higher than all helical pro-CPG2 controls, and for "high probability" non-inhibitors, the enrichment ratio at 30 μM methotrexate needed to be lower than that at 50 μM and 100 μM.

**Preparation of individual pro-CPG2 design constructs.** The vector backbone and CPG2 scaffold were amplified from the helical design plasmids by PCR (Table S10, primers 21-22). The sixteen designs were ordered as gBlocks (Integrated DNA Technologies Inc.) and inserted into this vector using Gibson assembly (6).

The TEV cleavage site was substituted for an MMP site using primer extension PCR (Table S10, primers 29-32) followed by QuikChange mutagenesis (Agilent Technologies Inc.).

Disulfide variants were generated using QuikChange Multi-Site Mutagenesis (Agilent Technologies Inc.) (Table S10, primers 23-28), starting from the Pro-CPG2-1 and Pro-CPG2-2 TEV variants.

**Preparation of pro-domain constructs.** The pro-domains were amplified using PCR (Table S10, primers 33-36) and re-inserted into pET15b, linearized by NdeI and BamHI.

**Protein expression.** Plasmid DNA was transformed into BL21(DE3) cells. LB media containing 100 μg/mL ampicillin was inoculated with a single colony and grown overnight at 37ºC. ZYP-5052 media (14) (5 mL for screening, 500 mL for other applications) was inoculated with the starter culture, using 1/50$^{th}$ of the culture volume, and grown for 3 hours at 37ºC followed by 18ºC overnight. The cells were harvested by centrifugation at 4ºC and re-suspended in 50 mM Tris, 300 mM NaCl, 30 mM imidazole, pH 7.4 supplemented with lysozyme, PMSF, and DNase I. For screening, 1.5X BugBuster (EMD Millipore) was also added.

**Protein purification for pro-CPG2 design activity screening.** The BugBuster cell suspension was incubated at room temperature for 20 minutes, and then 40 minutes at 4ºC. The lysate was centrifuged at 20,000×g, and the supernatant was applied to 150 μL of Ni-NTA agarose resin (Qiagen), shaking for 1 hour. The resin was washed with 50 mM Tris, 300 mM NaCl, 30 mM imidazole, pH 7.4, and bound protein was eluted using 50 mM Tris, 300 mM NaCl, 300 mM imidazole, pH 7.4. The protein purity was analyzed using SDS-PAGE, and the protein was dialyzed against 50 mM Tris, 100 mM NaCl, pH 7.4.

**Protein purification for Michaelis-Menten kinetics and structural biology.** The cells were lysed by sonication (Fisherbrand Model 705 Sonic Dismembrator), and the soluble fraction was isolated by centrifugation at 20,000×g for 30-60 minutes. After filtering, the supernatant was applied to a 5 mL NiNTA Superflow column (Qiagen) equilibrated with 50 mM Tris, 300 mM NaCl, 30 mM imidazole, pH 7.4 using an Äkta Pure (GE Life Sciences). The protein was eluted with a linear gradient of 30 mM – 300 mM imidazole, and fractions containing the CPG2 designs were pooled.

The samples were desalted into 50 mM Tris, 300 mM NaCl, pH 7.4 using a HiPrep 26/10 Desalting column (GE Life Sciences) and then supplemented with 2.5 mM $CaCl_2$ and ~1 unit of thrombin/mg protein. The sample was incubated at room temperature for 3 hours. Uncleaved protein and thrombin were removed by re-applying the protein to a 5 mL NiNTA Superflow column (Qiagen) with a 1 mL HiTrap Benzamidine Fast Flow (high sub) column (GE Life Sciences) connected in series. The flowthrough was concentrated to ~2 mL and applied to a HiLoad 16/600 Superdex 200 pg column (GE Life Sciences) equilibrated with 50 mM Tris, 100 mM NaCl, pH 7.4. The purity of the sample was assessed using SDS-PAGE, and fractions containing pure protein were pooled and concentrated to 10-20 mg/mL.

**Protein purification of the Isolated Pro-Domains.** The cells were lysed by sonication as described above. The supernatant was applied to 5 mL of Ni-NTA agarose resin (Qiagen), shaking for 1 hour. The resin was washed with 50 mM Tris, 300 mM NaCl, 30 mM imidazole, pH 7.4, and bound protein was eluted using 50 mM Tris, 300 mM NaCl, 300 mM imidazole, pH 7.4. The protein purity was analyzed using SDS-PAGE, and the protein was dialyzed against 20 mM sodium phosphate, 100 mM NaCl, pH 7.4.

**Pro-CPG2 proteolytic re-activation.** For the TEV protease variants, the protein was diluted to 1.0-1.5 mg/mL in 50 mM Tris, 100 mM NaCl, pH 7.4. TEV protease, stored at 1 mg/mL in 50 mM sodium phosphate, 200 mM NaCl, 10% glycerol, 2 mM EDTA, 10 mM DTT, pH 7.5, was added to a final concentration of 0.1 mg/mL. The samples were incubated at 4ºC for 48 hours. In some experiments, these samples were dialysed against 50 mM Tris, 100 mM NaCl, pH 7.4 using a 20,000 molecular weight cutoff membrane sufficient to retain the CPG2 protein while allowing passage of the "free" pro-domains following cleavage. Following proteolysis, the samples were tested for activity as described above. In addition, samples were analyzed using LC-MS with a Velos Mass Spectrometer (Thermo Scientific). In brief, the sample was loaded in a mixture of 85% buffer A (0.2% formic acid) and 15% buffer B (0.1% formic acid in acetonitrile). After 3 minutes, a linear gradient from 15%-100% buffer B was applied over 7 minutes, after which the column was held at 100% B for another 3 minutes. Data were analyzed using Thermo Scientific Protein Deconvolution software.

For the MMP Pro-CPG2 variants, the protein was diluted to 2 µM (approximately 100 µg/mL) in 50 mM Tris, 100 mM NaCl, 10 mM $CaCl_2$, 5 µM $ZnSO_4$, pH 7.4, with a final concentration of 40 nM MMP-2 (Sigma Aldrich, 10 µg in 10 mM sodium phosphate pH 7.5, 0.1 mM $CaCl_2$ reconstituted in 0.1% bovine serum albumin). The samples were incubated at 4ºC for 24 hours, and the samples were tested for activity as described above.

**Circular dichroism of individual pro-domains.** The pro-domain solutions were filtered (0.2 µm) and diluted to 0.1 mg/mL using 20 mM sodium phosphate, 100 mM NaCl, pH 7.4. CD spectra were acquired on an Aviv Model 420SF CD Spectrometer at 4°C in 1.0 mm cuvettes over the wavelength range of 190-260 nm in 2 nm increments with an averaging time interval of 2 s. Temperature-dependent CD spectra were recorded from 10ºC to 100ºC in 5ºC increments. Buffer and duplicate measurements were recorded periodically. The CD profiles were smoothed using the Savitzky-Golay filter (15), and the buffer spectrum at the appropriate temperature was subtracted from the sample spectra. The average baseline molar ellipticity between 250-260 nm was subtracted from the CD curve.

**Isothermal titration calorimetry (ITC).** Thermodynamic binding parameters for the association of $CPG2_{CP\text{-}N89}$-K177A with Pro-Domain-1 were determined calorimetrically employing a MicroCal VP-ITC (Malvern Panalytical, Northampton, MA). Protein ($CPG2_{CP\text{-}N89}$-K177A) and Pro-Domain-1 stock solutions were prepared and dialyzed exhaustively against 50 mM sodium phosphate, 100 mM NaCl, pH 7.4. Each ITC experiment consisted of thirty consecutive 10.0 µL injections during which the reaction heat is monitored and integrated over a 5.0 minute period under constant stirring conditions. The experimental protocol has been designed to ensure that there is a tenfold excess of Pro-Domain-1 in the titration syringe (i.e., 150, 250, or 500 µM) relative to the initial enzyme concentration (i.e., 15, 25, or 50 µM) in the sample cell. The resultant binding isotherms are generated by recording the integrated reaction heats normalized for peptide concentration versus the Pro-Domain:Enzyme ratio. Inspection of these profiles reveals that formation of the enzyme-Pro-Domain-1 complex is characterized by mid-low micromolar binding affinities and relatively low enthalpies. Accordingly, we have employed the NITPIC/SEDPHAT suite of software programs (16, 17) to facilitate unbiased baseline assignment and peak integration. A nonlinear least squares fit of the resultant profile to a single site binding model yields thermodynamic parameters for the $CPG2_{CP\text{-}N89}$-K177A:Pro-Domain-1 complex including the affinity ($K_a$), Gibbs free energy ($\Delta G$), enthalpy ($\Delta H$), and entropy ($\Delta S$).

**Small-angle X-ray scattering.** For SAXS, protein samples were prepared as described above. The protein was diluted to 10 mg/mL using 50 mM Tris, 100 mM NaCl, pH 7.4, and 2 mM methotrexate (if applicable). Note that methotrexate is hydrolyzed by CPG2 prior to data collection. These samples were microdialyzed against the same buffer to establish a "matching" buffer for SAXS buffer subtraction.

Following dialysis, the protein samples were diluted with their matched buffer by a factor of two and four, yielding protein concentrations of ~2.5 and ~5 mg/mL. SAXS data were collected for these samples, the ~10 mg/mL sample, and two matched buffer samples at the SIBYLS beamline (beamline 12.3.1) at the Advanced Light Source (18). An exposure of 10 seconds split over 32 frames was collected. The buffer signal was subtracted from the protein curves, and the concentration series was merged in PRIMUS (19) such that noisy data at higher q-ranges and interparticle repulsion at low q-ranges were eliminated from the merged scattering curves. Particle distance distribution plots were generated using GNOM (20), and ScÅtter (Rodic & Rambo, www.bioisis.net) was used to determine other SAXS parameters, including the particle density (21, 22), Porod exponent (22), and volume of correlation (23).

**X-ray Crystallography.** CPG2$_{CP\text{-}N89}$-K177A-MMP-Helix (best helical pro-domain sequence: AEAAWKEAEAKDWAAKA) was prepared as described above to a concentration of 19 mg/mL in 50 mM Tris 100 mM NaCl pH 7.4 and 0.2 $ZnSO_4$. Crystals were grown using hanging drop vapor diffusion. Reservoirs containing 750 µL of 20 mM Tris pH 8.0, 10% glycerol, and 10% PEG 3350 were prepared in 24 well hanging drop vapor diffusion plates. Equal volumes of protein solution and reservoir solution were mixed on a cover slip and suspended over the reservoir. The plate was incubated at 20ºC, and crystals were obtained in 1-2 weeks.

Crystals were harvested and flash-frozen in a nitrogen cryostream. Data were collected on a Rigaku MicroMax-007 generator equipped with an RAxis IV++ detector. Data were processed using xia2 (24) with the XDS and XSCALE (25) pipeline, indexing with peaks from all images and with the ice rings filter enabled. The structure was solved by molecular replacement using Phaser (26) and positional and B-factor refinement was performed using Refmac (27) with non-crystallographic symmetry constraints. Manual inspection and model building was performed with Coot (28).

#### SUPPLEMENTARY RESULTS

##### S1 Selection of designs for inclusion in the library

After completing the process of pro-enzyme designs using the small protein pro-domain set, we excluded designs with low Rosetta scores or repulsive energy scores, few contacts with the active site core region of CPG2, a high number of buried unsatisfied hydrogen bonds, and poor zinc-glutamate interaction geometry. At this point, we retained 15,418 designs (Table S4), of which 1668 were from the original design set, 8757 were from the fixed backbone design set, and 4993 were from the explicit hydrogen bond network set (Figure S4). Distributions of overall scores and specific score terms between the three sets were remarkably consistent (Figure S4).

Next, we eliminated designs that featured large numbers of mutations, poor shape complementarity between the pro-domain and scaffold, and low numbers of contacts between the pro-domain and critical interaction sites on the CPG2 scaffold. This filtering further reduced the number of designs to 7511, which was finally reduced to 7469 by removing the worst scoring designs, by Rosetta energy, from each design set.

Finally, we included “control” sequences for each of four helical pro-domain protein designs with known inhibitory properties (8 DNA sequences for each of three mildly inhibiting protein sequences, and 7 DNA sequences for one non-inhibiting protein sequence) to serve as internal standards for testing the designed enzyme pool. Three of these sequences were the pro-domains identified as being effective inhibitors, while the fourth was a pro-domain known to be an ineffective inhibitor. These controls completed our library of 7500 pro-domains, which included 7469 candidate designs and 31 control sequences that allow the relative activities determined in our high-throughput assay to be compared to pro-domains with known activities.

#### S2 Computational assessment of Pro-CPG2 design models

We assessed the overall quality of the computational models using a series of Rosetta simulations that probe specific characteristics (Figure S8). In the first (left panels), we attempted *ab initio* structure prediction of the pro-domains, in the absence of $CPG2_{CP\text{-}N89}$-K177A. All three pro-domains showed a folding funnel, with the lowest energy structures characterized by the lowest RMSDs. In the case of Pro-Domains-1 and -3, there appears to be a single, low energy species, with a particularly “narrow” funnel for Pro-Domain-1. Conversely, Pro-Domain-2 showed three low energy species, indicating that this structure may exhibit multiple folded states.

Our second analysis also performed *ab initio* structure prediction of the pro-domains, but this time in the context of the $CPG2_{CP\text{-}N89}$-K177A “scaffold” (Figure S8, middle panels, script 6 in Table S9). In contrast to the previous calculation, this simulation incorporates any role the scaffold might play in stabilizing the correct fold. Once again, folding funnels are observed, though they were shallower than in the previous case, and with Pro-CPG2-1 showing the most distinct folding funnel. All three pro-enzymes exhibit evidence of distinct higher energy folding funnels that mirrored the lowest energy funnel (Figure S8).

Finally, maintaining the structure of both the pro-domain and $CPG2_{CP\text{-}N89}$-K177A fixed, we sampled backbone conformations of the linker connecting the pro-domain to the scaffold, thereby simulating the binding and release of the fully folded pro-domain to the scaffold (Figure S8, right panels, script 7 in Table S9). In the case of Pro-CPG2-2 and -3, an extremely steep folding funnel is observed, reflective of a single, low energy binding mode and a large ensemble of unbound conformations. In contrast, Pro-CPG2-1 shows a comparatively flat energy landscape, with the lowest energy structures occurring with an elevated RMSD.

Taken together, the three sets of simulations reveal that Pro-CPG2-1, the most strongly inhibited pro-enzyme design has the most pronounced *ab initio* folding funnel. Our finding suggests that this characteristic is more important to the design of a strongly inhibiting pro-domain than conformational landscapes obtained by treating the pro-domain as a rigid body. Forward folding of the pro-domain thus may provide an important metric for evaluating and selecting higher quality designs *in silico*. In contrast, the rigid body sampling appears to have more limited usefulness in assessing design quality and may also reflect that the possibility of additional low-energy binding modes may not completely exclude strong inhibitory properties for a design.

#### S3 ITC reveals potential interdependence of the two pro-domain binding sites

The Pro-CPG2-1 design model allows for one Pro-Domain-1 molecule to bind each CPG2 protomer and does not suggest any direct interaction between the two Pro-Domain-1 molecules when both are bound. We might therefore expect that the two Pro-Domain-1 binding sites in the $CPG2_{CP\text{-}N89}$-K177A dimer are equivalent. Intriguingly, there is some evidence that the two sites are not equivalent, as our ITC data suggest that binding of the first Pro-Domain-1 may impact binding of the second. Unfortunately, the low ITC-derived reaction heats preclude a detailed analysis of this interdependence. Future studies employing complementary approaches may provide valuable insight into the CPG2:Pro-Domain-1 binding mechanism and, by extension, the enzyme mechanism of CPG2, with the potential to guide subsequent design efforts.

#### S4 Model-free SAXS data analysis

SAXS provides powerful tools for assessing protein flexibility (21, 22). Examination of the Kratky plots confirms that all of the variants analyzed using SAXS exhibited well-folded, globular structures. This trend was confirmed quantitatively using the Porod exponent (22), which ranged from 3.7-4.0 across all variants (Table S8). The protein density, as assessed using the Porod-Debye law (21), ranged from 0.86-1.28 g/mL (Table S8). In comparison to typical particle densities obtained using SAXS and the theoretical protein density of 1.35-1.37 g/mL (21), this indicates some degree of conformational flexibility, with the auto-inhibited variants showing lower density, and therefore greater flexibility, than $CPG2_{CP\text{-}N89}$-K177A. Furthermore, addition of hydrolyzed methotrexate triggered a substantial increase in particle density, from 1.19 g/mL to 1.28 g/mL, for $CPG2_{CP\text{-}N89}$-K177A, while the pro-enzyme designs had densities that were largely unchanged in comparison (Table S8).

The dimensionless Kratky plots (21, 22) also provide valuable information regarding the nature of particle flexibility (Figure 4CD). In the $R_g$-based plot (Figure 4C), if a peak occurs at $\sqrt{3}$ on the horizontal axis and 1.104 on the vertical axis, it is indicative of a compact, globular particle. If the peak is shifted up and to the right, this indicates that the particle is more elongated, and if the curve becomes a hyperbolic plateau, this is indicative of a disordered particle. $CPG2_{CP\text{-}N89}$-K177A and all of the pro-enzymes show clear peaks, indicative of a well-folded particle. In the absence of hydrolyzed methotrexate, the $CPG2_{CP\text{-}N89}$-K177A peak is substantially shifted up and to the right, suggestive of an elongated particle. Addition of hydrolyzed methotrexate causes the peak to shift back, suggesting that the addition of hydrolyzed methotrexate causes the enzyme to adopt a more compact conformation, consistent with a shift towards the “closed” conformation. The pro-enzyme forms are even further shifted, an effect which can be attributed to the pro-domain adding mass towards the center of mass of enzyme, resulting in a less elongated particle. The peaks in the pro-enzyme forms are largely unresponsive to addition of hydrolyzed methotrexate. These trends appear to be very subtly even more pronounced in the Pro-CPG2-1-Disulfide variant, as compared to Pro-CPG2-1.

The $V_c$-based dimensionless Kratky plot (Figure 4D) provides insight into the surface-to-volume ratio, with a peak at $\sqrt{3}$ on the horizontal axis with a height of 0.82 indicating a surface-to-volume ratio consistent with a perfect sphere, and a downwards shift indicating a larger surface-to-volume ratio. The peak heights in this plot are in a fairly narrow band, suggesting that the exposed surface area is fairly similar in all cases. $CPG2_{CP-N89}$-K177A in the absence of hydrolyzed methotrexate has the highest surface-to-volume ratio, followed by $CPG2_{CP-N89}$-K177A in the presence of hydrolyzed methotrexate and the pro-enzymes in the absence of hydrolyzed methotrexate, consistent with the burial of surface area through the closure of the enzyme and/or the binding of the pro-domain to the native CPG2 surface. The lowest surface-to-volume ratio was obtained for the pro-enzyme in the presence of hydrolyzed methotrexate.

It should also be noted, particularly when examining the $R_g$-based Kratky plots (Figure 4C), that the enzyme in the absence of hydrolyzed methotrexate shows a somewhat bimodal peak shape, with a peak at around 2 and another peak at approximately 3.5. This bimodal character is diminished in the presence of hydrolyzed methotrexate, and particularly in the case of the Pro-CPG2-1-Disulfide variant. The bimodal peaks may suggest two structural populations, perhaps corresponding to the “open” and “closed” conformations.

#### SUPPLEMENTARY FIGURES

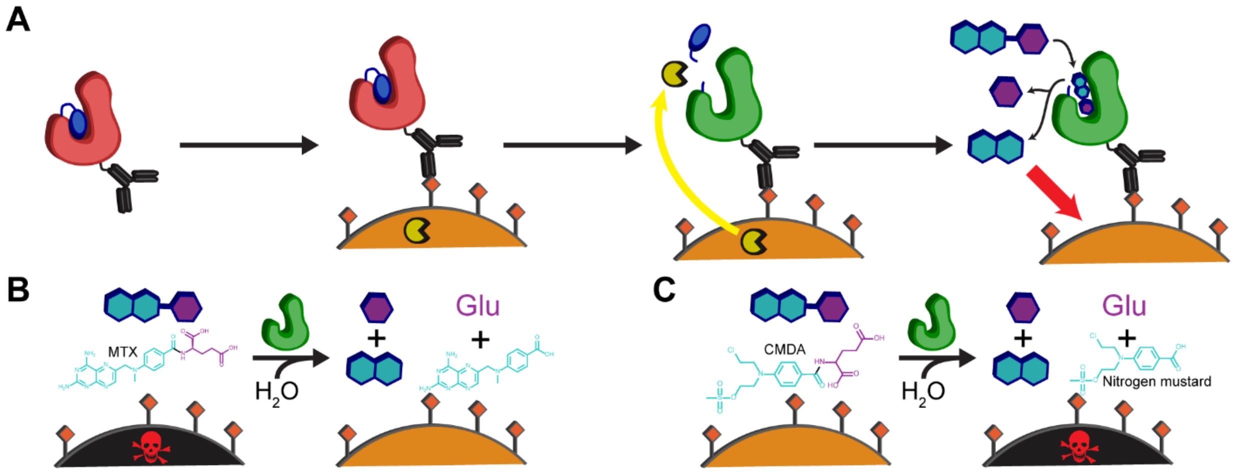

**Figure S1.** Utility of a protease-activatable form of CPG2. (A) Mechanism of activation of antibody-linked Pro-CPG2. The antibody (black) directs the inactive enzyme (red) to a protease-expressing cell (cell is orange, protease is yellow, antibody's targets are orange diamonds). The protease (yellow) cleaves the linker that connects the pro-domain (blue) to the enzyme, activating it (green). The enzyme can then process small molecules (teal and purple) which can have an effect (red arrow) on the protease-expressing cell. (B) Activated CPG2 can cleave and detoxify the drug, methotrexate, rescuing the protease expressing cells (black to orange). (C) Activated CPG2 can cleave and activate the nitrogen mustard prodrug CMDA, killing the protease expressing cells (orange to black).

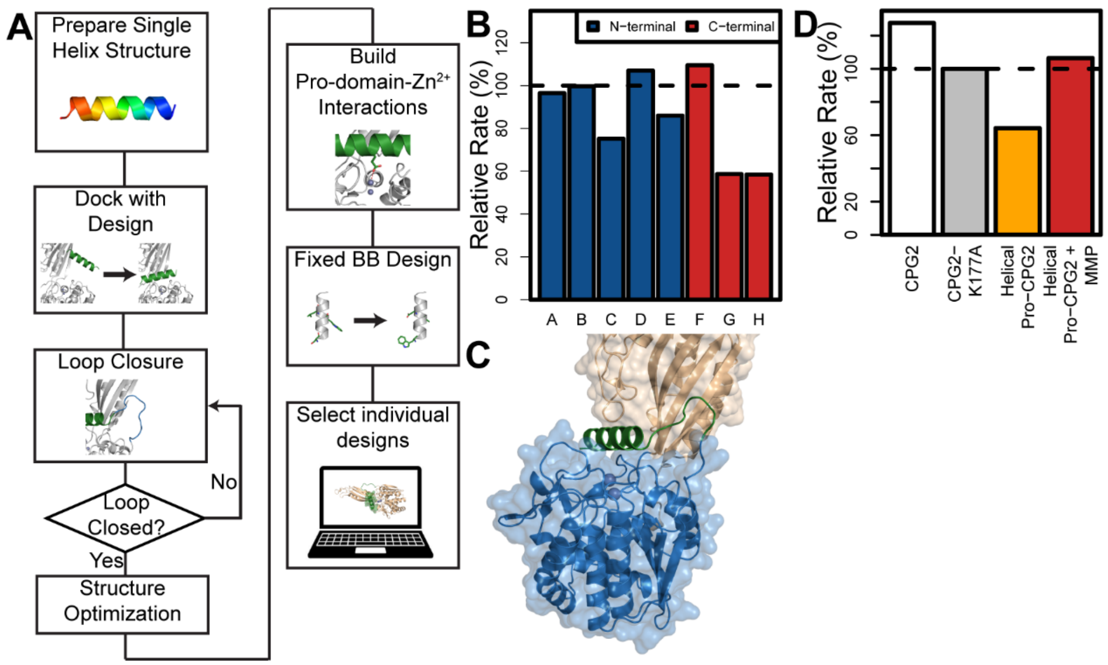

**Figure S2.** First-generation, helical design strategy. (A) Flow chart depicting the design strategy for the first-generation, helical design. (B) Preliminary relative molar specific activity data comparing the activity of eight helical pro-enzyme designs (designated A-H, see also Table S1 and Table S10) to the activity of $CPG2_{CP\text{-}N89}$, including any mutations present in the pro-enzyme designs. The terminus to which the pro-domain is attached (N for variants A-E and C for variants F-H) for each pro-enzyme is highlighted. (C) Model of one of the first-generation, helical designs, highlighting the pro-domain (green), the catalytic domain (blue), and the dimerization domain (wheat). (D) The relative molar specific activity of the first-generation helical design before (orange) and after (red) incubation with the appropriate protease to remove the pro-domain. Molar specific activities are given relative to the circular permutant, $CPG2_{CP\text{-}N89}$-K177A. The wild-type circular permutant, $CPG2_{CP\text{-}N89}$, is also shown in panel D.

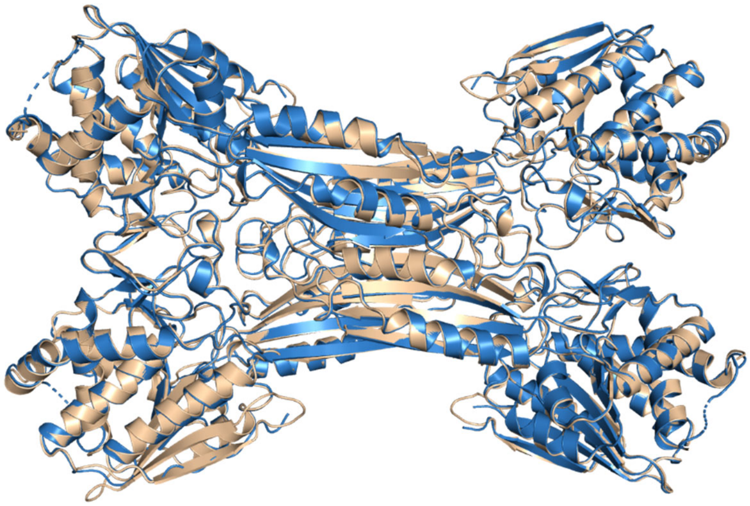

**Figure S3.** Crystal structure of helical pro-CPG2 design. The individual chains of PDB ID 7M6U (blue) are overlaid onto the individual chains of CPG2 (PDB ID 1CG2, wheat). The Cα RMSD range from 0.26-0.52 Å when comparing all 1CG2 chains to all 7M6U chains.

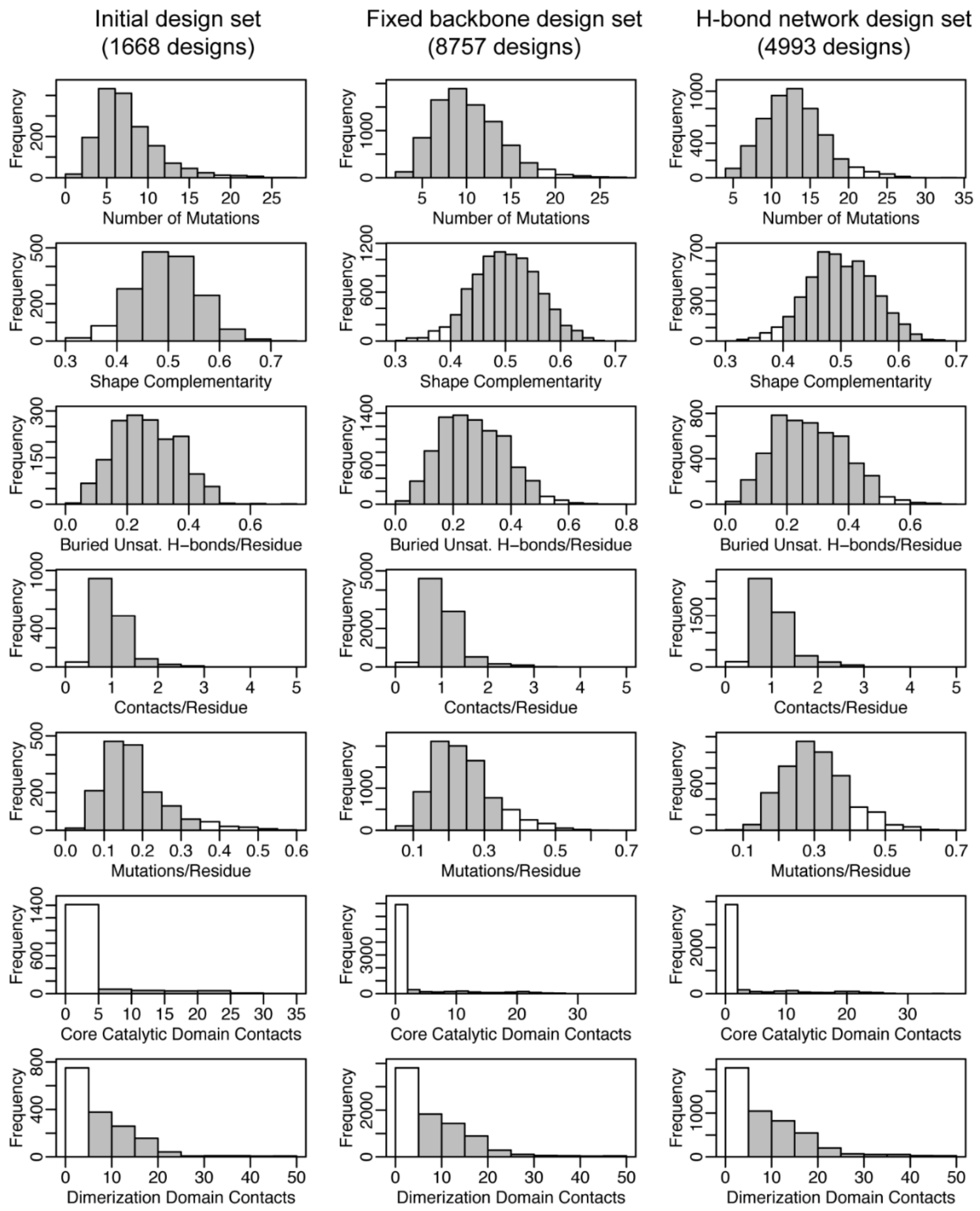

**Figure S4.** Distribution of computational scores in each of the three design sets. Shaded bars were included in the design library. Note that a shaded design from one metric could be excluded by falling outside of the shaded region for another metric. Number of mutations is the total number of mutations introduced. Shape complementarity is calculated between the CPG2 scaffold and pro-domain. Buried unsatisfied H-bonds are calculated across the scaffold-pro-domain interface. Core catalytic domain contacts and dimerization domain contacts include sidechain carbon-carbon contacts between the pro-domain and residues 198-211, 358-362, and 375-383 and residues 215-219 and 320-324, respectively (wild-type CPG2 numbering). Per residue metrics are normalized to the number of pro-domain residues in the design.

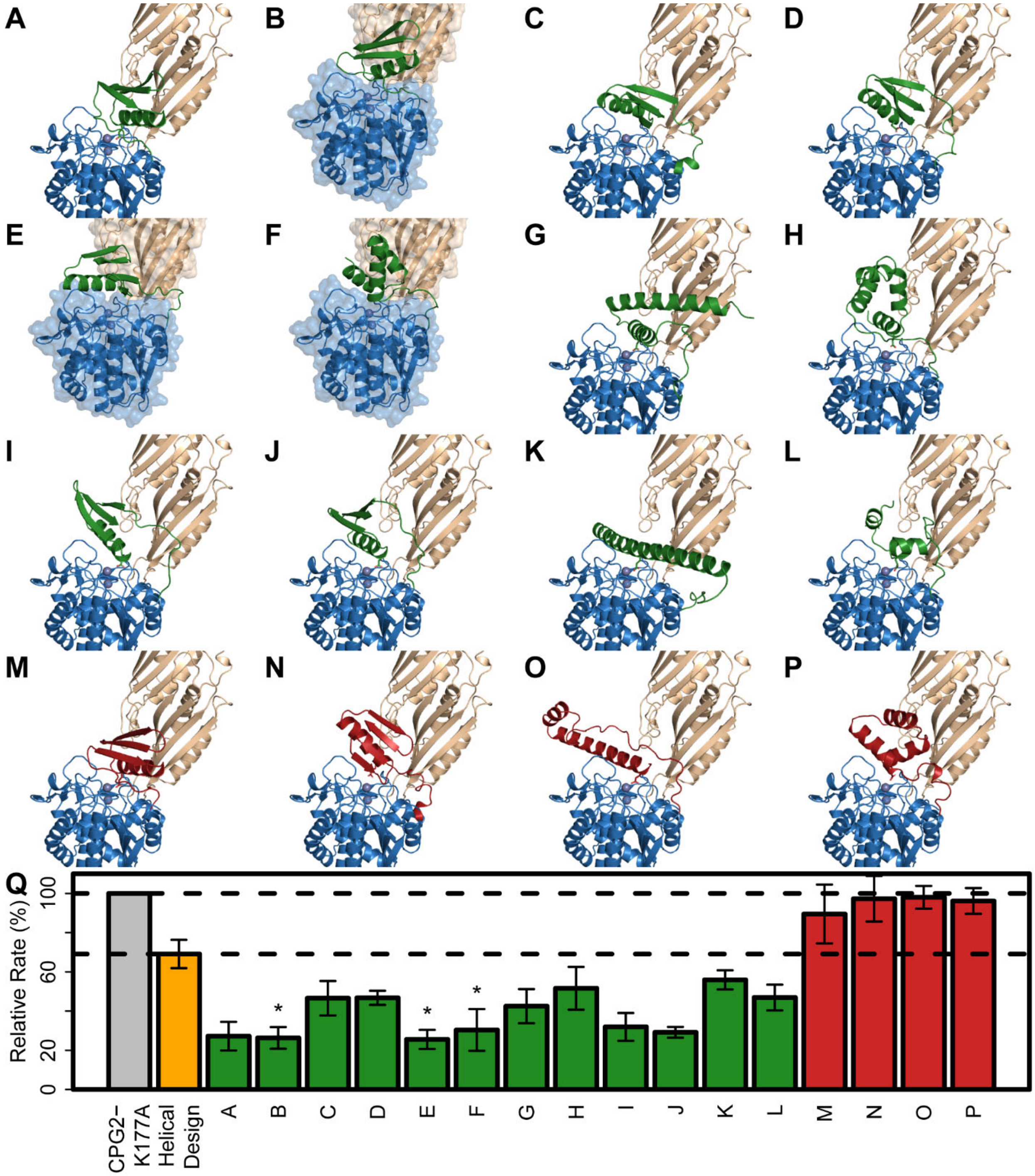

**Figure S5.** Rosetta models of the 16 designs selected for individual expression. (A-L) The designs predicted to have strongly inhibiting pro-domains (green). (M-P) The designs predicted to have poorly inhibiting pro-domains (red). The native CPG2 structure is shown in blue (catalytic domain) and wheat (dimerization domain). Pro-CPG2-1, -2, and -3 have a translucent surface (blue and wheat) shown for the native structure. (Q) Relative molar specific activity data for $CPG2_{CP\text{-}N89}$-K177A, the original "helical" pro-domain design, and the 16 designs shown in panels A-P. Dashed lines indicate the activity levels for $CPG2_{CP\text{-}N89}$-K177A and the helical pro-CPG2. The designs selected for detailed characterization are denoted by an asterisk.

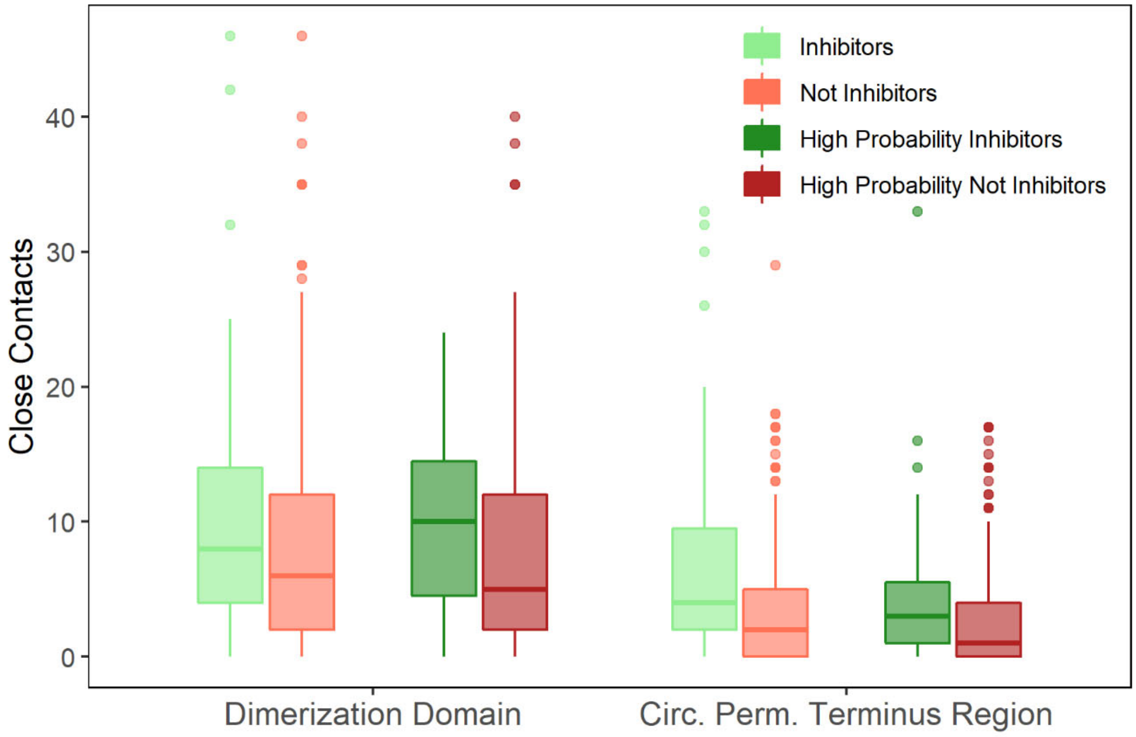

**Figure S6.** Comparison of close contact count distributions. Comparison is made between the pro-domain and selected dimerization domain (residues 215-219 and 320-324) and catalytic domain (residues 173-182) residues. Contact counts are computed using Rosetta's AtomicContactCount filter. Sidechain carbon-carbon contacts between the designed pro-domain and the specified CPG2 residues (wild-type CPG2 numbering) within 4.5 Å are counted. Predictions are made based on NGS enrichments. See also Table S6.

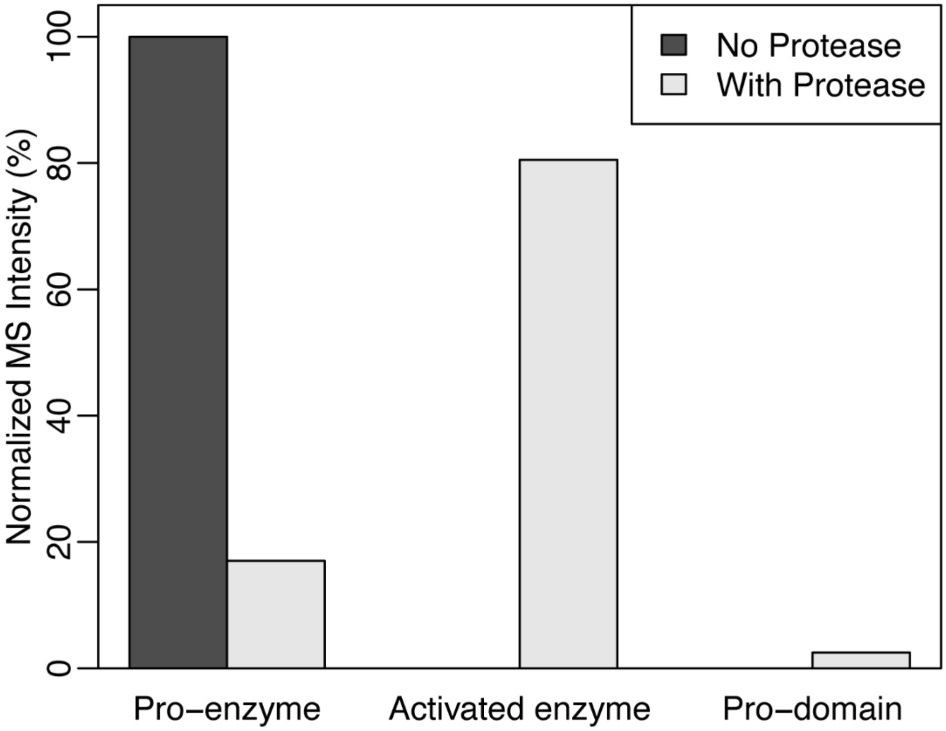

**Figure S7.** Mass spectrometry analysis of Pro-CPG2-1 before and after proteolytic cleavage. The relative intensity of the three components of Pro-CPG2-1, the uncleaved pro-enzyme, the cleaved, activated enzyme, and the cleaved pro-domain, are shown. Intensities are normalized to the total intensity of the three components.

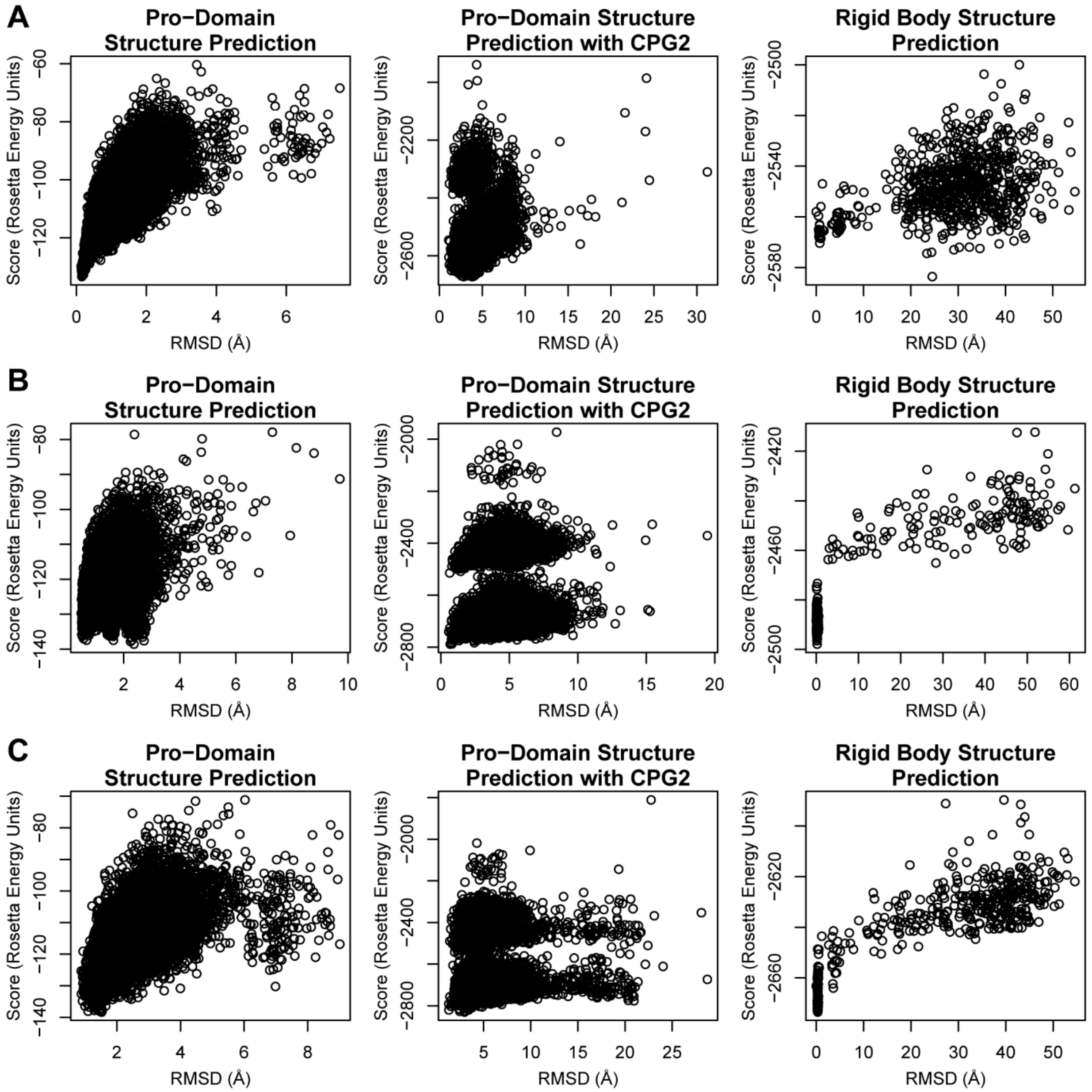

**Figure S8.** Computational assessment of the model quality of (A) Pro-CPG2-1, (B) Pro-CPG2-2, and (C) Pro-CPG2-3. The leftmost panels show the results of *ab initio* structure prediction of the pro-domains in isolation (10,000 decoys). The middle panels show the *ab initio* structure prediction of the pro-domains in the context of CPG2 structure (at least 10,000 decoys). The rightmost panels show the results of sampling CPG2-pro-domain linker conformations, allowing rigid body sampling of the enzyme-pro-domain conformation (750 decoys). It should be noted that the pro-domain is only superimposed on the reference structure in the leftmost panels (*ab initio* prediction in isolation). In the middle panels, strong outliers with energies greater than -1800 Rosetta Energy Units (up to 32 decoys, up to 0.3% of all decoys) are omitted for clarity.

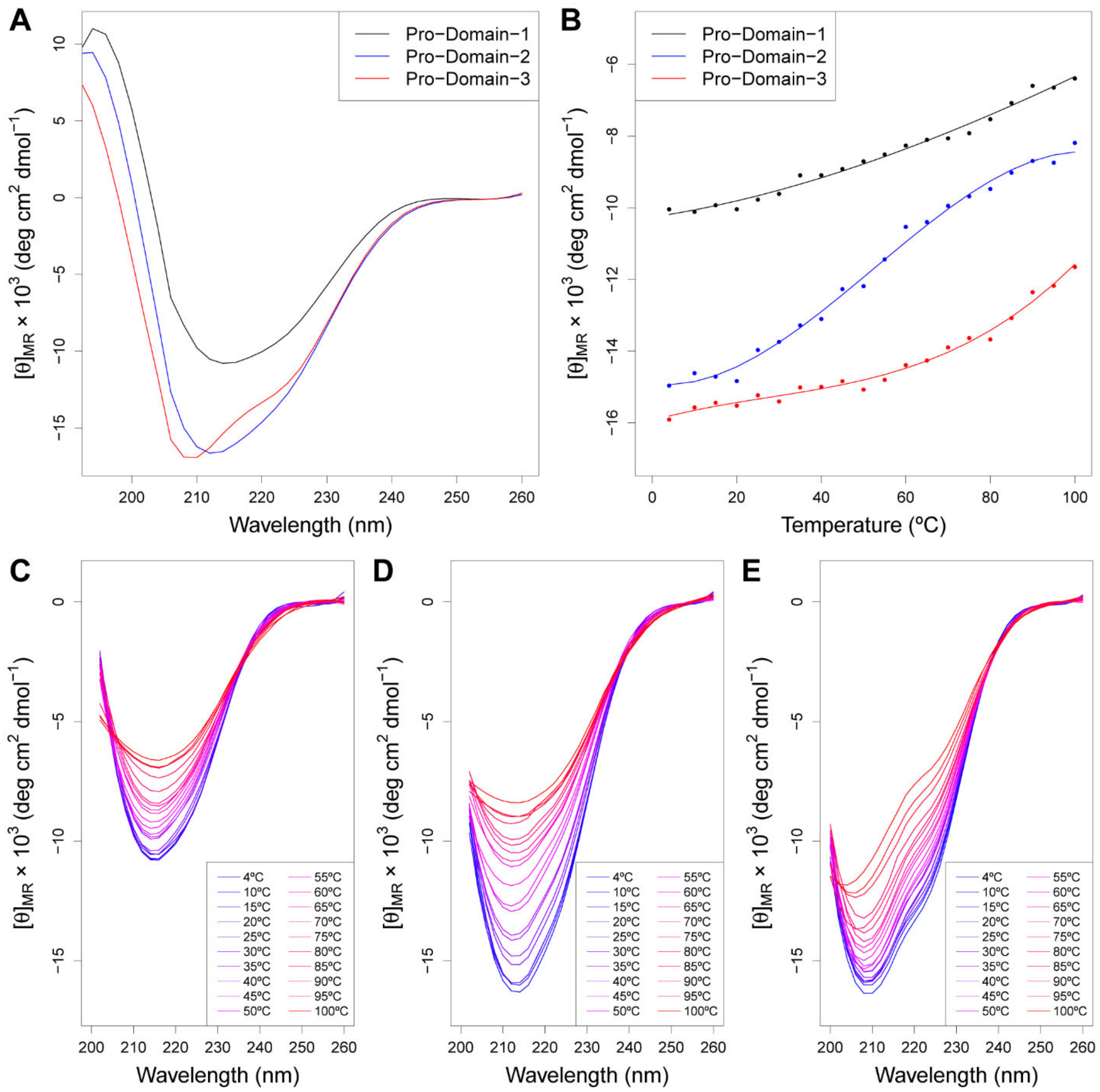

**Figure S9.** Circular dichroism spectra for the three Pro-Domains and their melting curves. (A) CD spectra of the three Pro-Domains at 4ºC. (B) Melting curves for the three Pro-Domains, recorded at the minimum wavelength of the CD spectra at 4ºC: 220 nm for Pro-Domain-1, 208 nm for Pro-Domain-2, and 206 nm for Pro-Domain-3. (C-E) The melting curves of (C) Pro-Domain-1, (D) Pro-Domain-2, and (E) Pro-Domain-3 shown as complete CD spectra. In all panels, CD spectra were baseline corrected and then smoothed using the Savitzky-Golay filter.

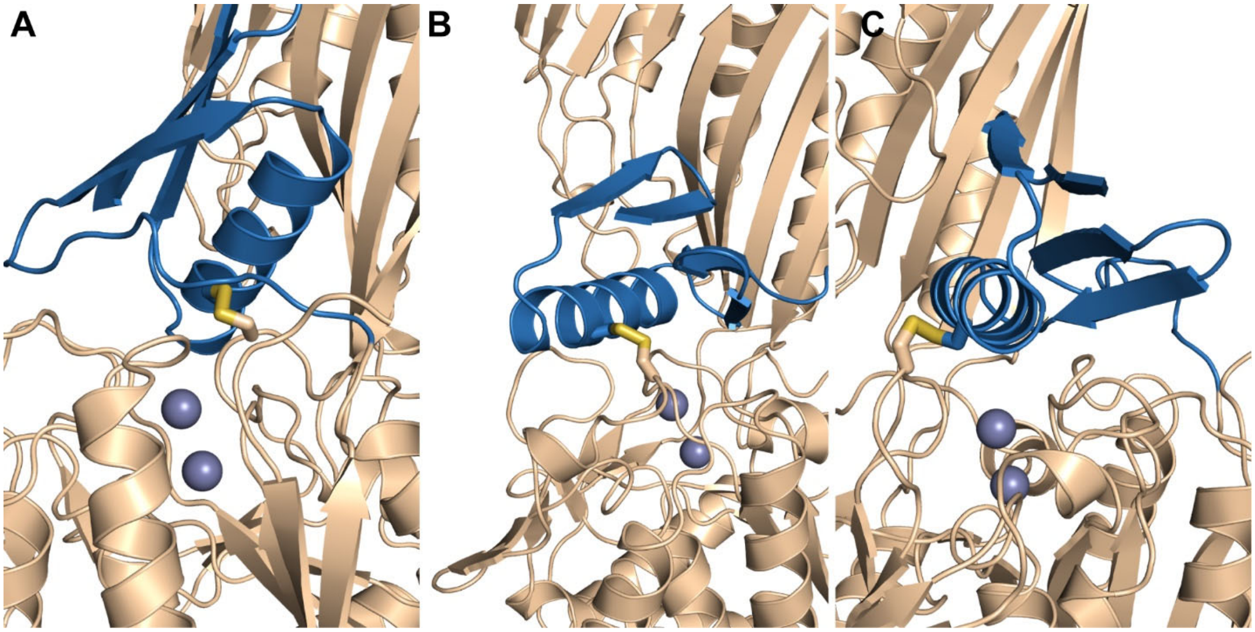

**Figure S10.** Rosetta models of the three disulfide designs. The CPG2 scaffold (wheat) and pro-domains (blue) are shown, with the connecting disulfide bond shown in sticks representation. The active site zincs are shown as grey spheres. (A) Pro-CPG2-1-Disulfide-1. (B) Pro-CPG2-2-Disulfide-2. (C) Pro-CPG2-2-Disulfide-3. See also Figure 3 and Table S5.

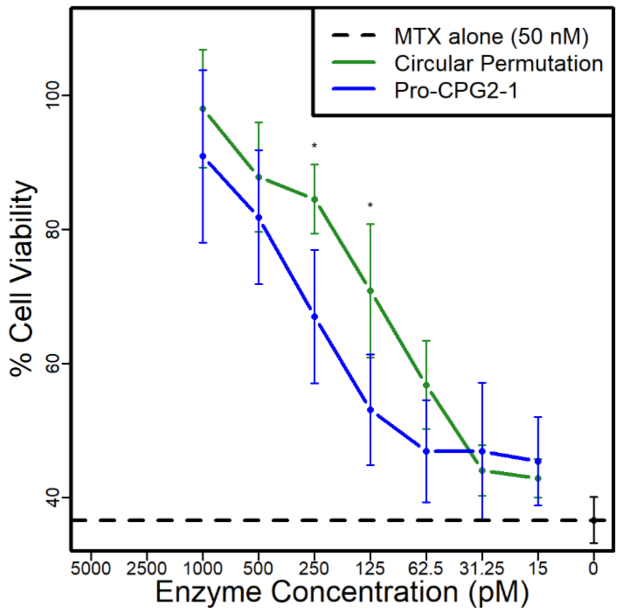

**Figure S11.** Pro-CPG2-1-Disulfide has reduced ability to cleave and detoxify methotrexate (MTX) compared to CPG2$_{CP-N89}$-K177A. PC3 cells were treated with MTX at 50 nM, and then treated with either CPG2$_{CP-N89}$-K177A or Pro-CPG2-1. Enzyme concentrations between 125-250 pM showed a qualitative difference (unpaired t-test, $p<0.1$) in PC3 cell viability between the two enzyme treatment groups, with cells treated with Pro-CPG2-1 exhibiting a lower survival rate than cells treated with active CPG2. The dashed line indicates the cell viability in the presence of MTX alone (no enzyme treatment). Error bars indicate the standard deviation. *= $p< 0.1$. Two-factor analysis of variance $p < 0.01$. See also Figure 5.

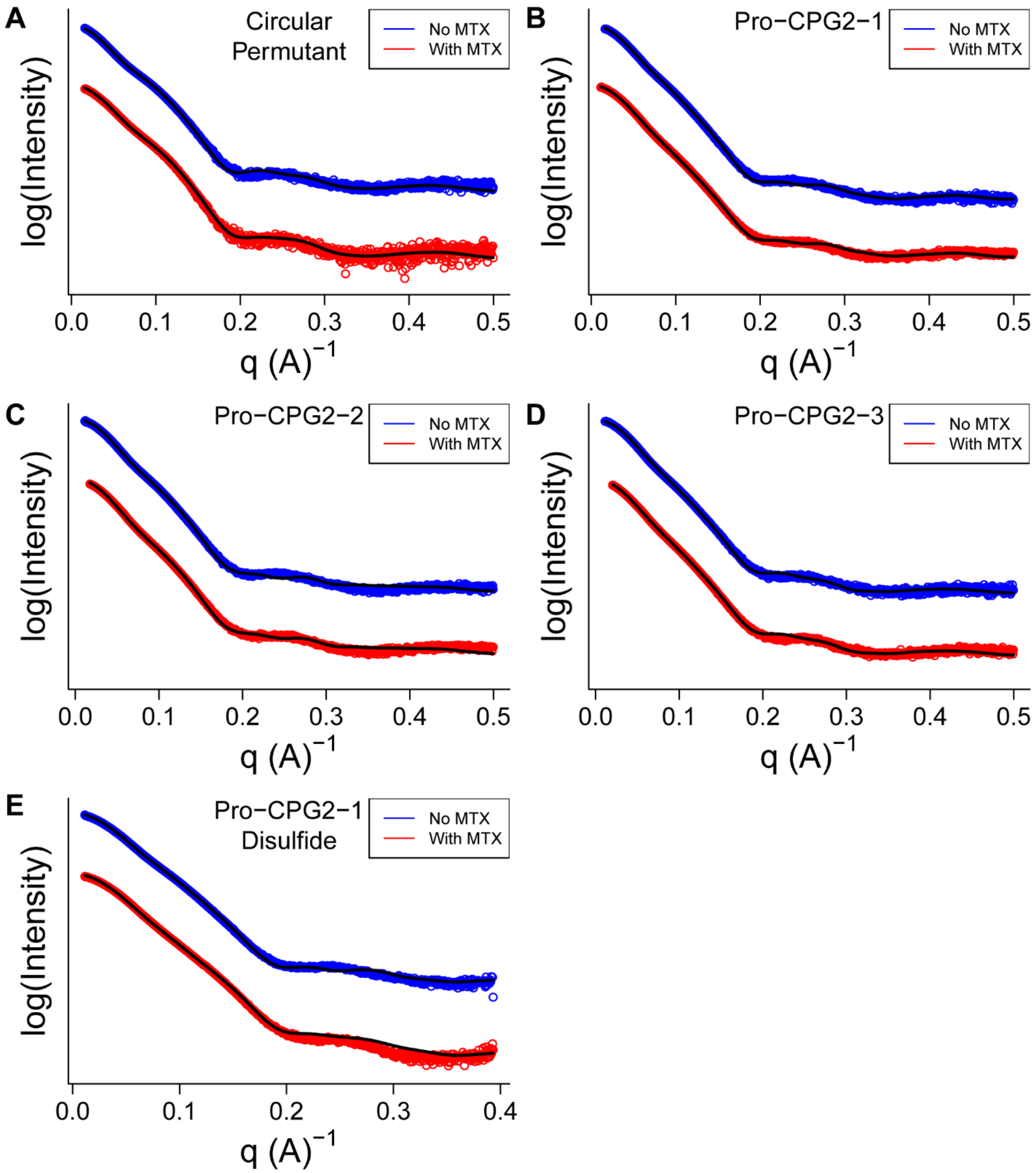

**Figure S12.** SAXS data and the two-state model fits for circularly permutated CPG2 and the pro-domains. (A-E) The raw experimental data for the samples with (red) and without (blue) the product, hydrolyzed methotrexate, along with the data fits (black line).

#### SUPPLEMENTARY TABLES

**Table S1.** Sequences of experimentally-characterized Pro-domains. All variants except Pro-Domain-1-Disulfide-1 include the K177A mutation (wild-type CPG2 numbering).

| Pro-Domain | Sequence |
|---|---|
| Helix G | AEAAWKEWEAKDWAAKA |
| Helix H | AEQAWKEWEAKDWAVKA |
| "Best" Helix | AEAAWKEAEAKDWAAKA |
| Pro-Domain-1 | QITYHVTGEEAARKIELMLAEANLEVEIHVDGDRYVLHAK |
| Pro-Domain-2 | STVHIGNETYYFPDSEEALKAAQAIAKKYGLTVERHGDTTTVE |
| Pro-Domain-3 | LRLAELIARKLAKNNQSPKAIAETLAKLGIDAEELKAALKKLT |
| Pro-Domain-1-Disulfide-1† | QITYHVTGEEAARKIELCLAEANLEVEIHVDGDRYVLHAK |
| Pro-Domain-2-Disulfide-2‡ | STVHIGNETYYFPDSEEALKAAQACAKKYGLTVERHGDTTTVE |
| Pro-Domain-2-Disulfide-3§ | STVHIGNETYYFPDSEEALKAAQCIAKKYGLTVERHGDTTTVE |

†Includes the K177C mutation (wild-type CPG2 numbering).
‡Includes the L118C mutation (wild-type CPG2 numbering).
§Includes the D387C mutation (wild-type CPG2 numbering).

**Table S2.** Crystallographic data for the CPG2$_{CP\text{-}N89}$-K177A-MMP-Helical Pro-domain Crystal Structure (PDB ID 7M6U).

| Data collection statistics | |
|---|---|
| Space group | $P2_1$ |
| Number of Molecules per Asymmetric Unit | 4 |
| a (Å) | 74.54 |
| b (Å) | 106.36 |
| c (Å) | 121.93 |
| α = γ (º) | 90.00 |
| β (º) | 107.48 |
| Wavelength (Å) | 1.54 |
| Resolution Range (Å)† | 39.27 – 2.59 (2.63 – 2.59) |
| Completeness (%)† | 97.20 (95.77) |
| Redundancy† | 2.86 (2.48) |
| I/σ | 6.81 (0.87) |
| $R_{merge}$ (%)† | 10.2 (94.6) |
| $R_{meas}$ (%)† | 12.5 (119.5) |
| $R_{pim}$ (%)† | 7.1 (71.9) |
| $CC_{1/2}$† | 0.952 (0.388) |
| **Refinement statistics** | |
| Total number of reflections (reflections in R-free | 45,894 (2346) |
| $R_{factor}$ (%) | 0.224 |
| $R_{free}$ (5% free test set) (%) | 0.279 |
| Number of atoms | 11,105 |
| Protein | 10,948 |
| Water | 132 |
| Catalytic $Zn^{2+}$ | 8 |
| Buffer components | 17 |
| RMSD | |
| Bond length (Å) | 0.007 |
| Bond angle (º) | 1.434 |
| Average atomic B-Factor (Å$^2$) | 60.0 |
| Protein (Å$^2$) | 60.2 |
| Water (Å$^2$) | 38.2 |
| Catalytic $Zn^{2+}$ (Å$^2$) | 48.2 |
| Buffer components (Å$^2$) | 88.5 |
| Ramachandran Plot | 1510 |
| Residues in Favoured Positions | 1444 |
| Residues in Allowed Positions | 61 |
| Residues in Disallowed Positions | 5 |

†Items in parentheses refer to the highest resolution shell.

**Table S3.** Computational metrics used to filter pro-CPG2 design candidates.

| Computational Metric | Threshold for removal from dataset |
|---|---|
| ShapeComplementarity (29) | < 0.4 |
| Buried unsatisfied hydrogen bonds/Pro-domain residue | > 0.5 |
| Percentage of pro-domain residues in contact with $CPG2_{CP-N89}$ | < 45% |
| Number of residues in contact with the dimerization domain beta-sheet region or cleft in the catalytic domain | < 1 |
| Number of mutated residues | > 16 (> 19 for HBNet designs) |
| Percentage of mutated residues | > 35% (> 40% for HBNet designs) |

**Table S4.** Summary of next-generation sequencing statistics.

| Experimental Stage | Number of sequences | Fraction of computational library (%) | Fraction of experimental library (%) | Fraction of high quality sequences (%) |
|---|---|---|---|---|
| Computational Library | 15,418 | 100 | N/A | N/A |
| Experimental Library | 7469 | 48 | 100 | N/A |
| High Quality Sequences | 1384 | 9.0 | 19 | 100 |
| Predicted Pro-CPG2s | 107 | 0.7 | 1.4 | 7.7 |
| High Probability Pro-CPG2s | 27 | 0.2 | 0.4 | 2.0 |
| Predicted Non-inhibitors | 515 | 3.3 | 6.9 | 37.2 |
| High Probability Non-inhibitors | 356 | 2.3 | 4.8 | 25.7 |
| Individually Expressed Pro-CPG2s | 12 | 0.08 | 0.16 | 0.9 |

**Table S5.** Michaelis-Menten parameters for the circular permutation of CPG2, the three Pro-CPG2 variants, and the disulfide variants.

| CPG2 Variant | $K_M$ (μM) | $k_{cat}$ ($s^{-1}$) | $k_{cat}/K_M$ ($s^{-1}$ $M^{-1}$) |
|---|---|---|---|
| $CPG2_{CP-N89}$-K177A | 50±6 | 109±5 | (2.2±0.3) × $10^6$ |
| Pro-CPG2-1 | 100±12 | 42±3 | (4.3±0.6) × $10^5$ |
| Pro-CPG2-2 | 136±19 | 60±5 | (4.4±0.7) × $10^5$ |
| Pro-CPG2-3 | 32±2 | 44.7±0.9 | (1.41±0.10) × $10^6$ |
| Pro-CPG2-1-Disulfide-1 | 163±19 | 24.6±1.6 | (1.5±0.2) × $10^5$ |
| Pro-CPG2-2-Disulfide-2 | 590±60 | 40±3 | (6.7±0.9) × $10^4$ |
| Pro-CPG2-2-Disulfide-3 | 160±30 | 6.6±0.6 | (4.3±0.8) × $10^4$ |

**Table S6.** Differences in computationally computed metrics between predicted inhibited and predicted non-inhibited designs. Predictions are made based on NGS enrichments. p-values are computed using unpaired, two-sample t-tests. See also Figure S6.

| Computational Metric | Predicted Designs (107 inhibited, 515 not inhibited) | | | | "High probability" predicted designs (27 inhibited, 356 not inhibited) | | | |
|---|---|---|---|---|---|---|---|---|
| | p-Value | Mean (inhibited) | Mean (not inhibited) | ΔMean | p-Value | Mean (inhibited) | Mean (not inhibited) | ΔMean |
| Close contacts† (dimerization domain, residues 215-219 and 320-324) | 0.028 | 9.9 | 8.2 | 1.8 | 0.046 | 10.1 | 7.4 | 2.6 |
| Close contacts† (catalytic domain, residues 173-182) | $1.4 \times 10^{-5}$ | 6.4 | 3.3 | 3.1 | 0.103 | 5.2 | 2.9 | 2.3 |
| Rosetta‡ fa_atr | 0.00011 | -4748.4 | -4732.1 | -16.3 | 0.011 | -4751.2 | -4728.1 | -23.1 |
| Rosetta‡ fa_intra_rep | 0.0013 | 8.5 | 8.4 | 0.05 | 0.041 | 8.5 | 8.4 | 0.1 |
| Rosetta‡ hbond_bb_sc | 0.0035 | -144.3 | -143.2 | -1.1 | 0.011 | -145.0 | -143.2 | -1.9 |
| Rosetta‡ hbond_sc | 0.021 | -142.9 | -144.0 | 1.1 | 0.015 | -142.3 | -144.4 | 2.1 |
| Rosetta‡ p_aa_pp | 0.0019 | -200.0 | -198.8 | -1.2 | 0.010 | -200.7 | -198.9 | -1.8 |
| Rosetta‡ ref | $8.35 \times 10^{-13}$ | 292.6 | 285.5 | 7.1 | $1.97 \times 10^{-8}$ | 293.8 | 284.9 | 8.9 |

†Close contacts are computed using Rosetta's AtomicContactCount filter. Sidechain carbon-carbon contacts between the designed pro-domain and the specified CPG2 residues (wild-type CPG2 numbering) within 4.5 Å are counted.
‡Rosetta scoring terms that form part of the Rosetta ref2015 score. For full details, consult the Rosetta documentation at https://www.rosettacommons.org/docs/latest/Home.

**Table S7.** Summary of the thermodynamic binding profiles for the $CPG2_{CP-N89}$-K177A: Pro-Domain-1 complex.

| T (ºC) | $K_a \times 10^5$ ($M^{-1}$) | $K_d$ (μM) | ΔG (kcal $mol^{-1}$) | ΔH (kcal $mol^{-1}$) | TΔS (kcal $mol^{-1}$) |
|---|---|---|---|---|---|
| 5.0 | 1.5 ± 0.07 | 6.7 ± 0.30 | -6.6 ± 0.03 | 1.5 ± 0.21 | 8.1 ± 0.18 |
| 15.0 | 3.6 ± 0.40 | 2.8 ± 0.28 | -7.5 ± 0.30 | 0.6 ± 0.06 | 8.2 ± 0.40 |

**Table S8.** SAXS data parameters.

| CPG2 Variant | Guinier $R_g$ (Å)† | Real space $R_g$ (Å)† | $D_{max}$ (Å)† | Total† | Porod Volume (Å$^3$)‡ | Porod Exponent‡ | Volume of correlation (Å$^2$)‡ | Density (g/cm$^3$)‡ | SASBDB ID |
|---|---|---|---|---|---|---|---|---|---|
| CPG2$_{CP-N89}$-K177A | 38.7 | 38.89 | 140 | 0.64 | 117,701 | 3.8 | 537 | 1.19 | SASDLD2 |
| CPG2$_{CP-N89}$-K177A + MTX | 35.79 | 35.95 | 127 | 0.64 | 109,540 | 4 | 523 | 1.28 | SASDLE2 |
| Pro-CPG2-1 | 36.61 | 36.73 | 124 | 0.72 | 143,112 | 3.7 | 581 | 1.11 | SASDLF2 |
| Pro-CPG2-1 + MTX | 35.67 | 35.77 | 122 | 0.76 | 142,819 | 3.8 | 582.3 | 1.12 | SASDLG2 |
| Pro-CPG2-2 | 37 | 37.09 | 125 | 0.76 | 142,429 | 3.9 | 593.2 | 1.12 | SASDLH2 |
| Pro-CPG2-2 + MTX | 35.79 | 35.88 | 121 | 0.7 | 139,605 | 3.9 | 583.7 | 1.15 | SASDLJ2 |
| Pro-CPG2-3 | 37.13 | 37.24 | 130 | 0.7 | 143,847 | 4 | 604.2 | 1.11 | SASDLK2 |
| Pro-CPG2-3 + MTX | 35.81 | 35.92 | 123 | 0.73 | 138,247 | 4 | 584.1 | 1.16 | SASDLL2 |
| Pro-CPG2-1-Disulfide-1 | 36.17 | 36.29 | 126 | 0.65 | 137,590 | 3.9 | 575.2 | 1.16 | SASDLM2 |
| Pro-CPG2-1-Disulfide-1 + MTX | 35.13 | 35.2 | 123 | 0.66 | 145,330 | 3.7 | 582 | 1.10 | SASDLN2 |

†Calculated using GNOM (20).

‡Calculated using ScÅtter (Rodic & Rambo, www.bioisis.net).

**Table S9.** rosetta_scripts referred to in the text.

| Script Number | Path to Script | rosetta_scripts_scripts git repository SHA of initial commit |
|---|---|---|
| 1 | scripts/public/enzymedesign/proenzyme_design/ cpg2_proenzyme_dock_and_design_helix.xml | a8c1fa8 |
| 2 | scripts/public/enzymedesign/proenzyme_design/ cpg2_proenzyme_repack_matches_after_helix_ docking.xml | a8c1fa8 |
| 3 | scripts/public/enzymedesign/proenzyme_design/ cpg2_proenzyme_small_domain_glu_placestub. xml | a8c1fa8 |
| 4 | scripts/public/enzymedesign/proenzyme_design/ cpg2_proenzyme_prodomain_fixedbbdesign.xml | faf167a |
| 5 | scripts/public/enzymedesign/proenzyme_design/ cpg2_proenzyme_prodomain_hbnetdesign.xml | faf167a |
| 6 | scripts/public/enzymedesign/proenzyme_design/ abinitio_topbroker.xml | f5a5ff3 |
| 7 | scripts/public/enzymedesign/proenzyme_design/ cpg2_proenzyme_prodomain_energylandscape. xml | 7320921 |
| 8 | scripts/public/enzymedesign/proenzyme_design/ cpg2_proenzyme_disulfidize.xml | 8222e68 |

**Table S10.** Primers used in molecular biology. Residue numbers for CPG2 scaffold mutations are given in wild-type CPG2 residue numbering. Residue numbers for pro-domain residues are given assuming that the GS-TEV-GS linker spans residues 397-407 and the first residue of the pro-domain begins at residue 408.

| Primer Number | Primer Name | Primer Sequence |
|---|---|---|
| 1 | CPG2-K177A[†] | 5'-CAGATGAAGAAGCGGGTTCATTTGGTTCACGTGATTTGATCC-3' |
| 2 | CPG2$_{\text{CP-N89}}$-AddTEV-CHis-F | 5'-GGTTTTACAGTAACACGATCAAAATCAGCAGGTGGTAGCGAAAATCTTTATTTTCAAGGGGG-3' |
| 3 | CPG2$_{\text{CP-N89}}$-AddTEV-CHis-R | 5'-GGTGGTGGTGGTGGTGCTCGAGAGACCCCCCTTGAAAATAAAGATTTTCGC-3' |
| 4 | CPG2$_{\text{CP-N89}}$-TEV-CHis-Linearization-F | 5'-CTCGAGCACCACCACCACCACCACTGAGATC-3' |
| 5 | CPG2$_{\text{CP-N89}}$-TEV-CHis-Linearization-R | 5'-AGACCCCCCTTGAAAATAAAGATTTTCGCTACCACCTGC-3' |
| 6 | Helix F-F | 5'-GCGAAAATCTTTATTTTCAAGGGGGGTCTGCAGAAGCCCAAGCGAAAGAAGCCGAGG-3' |
| 7 | Helix F-R | 5'-GGTGGTGGTGGTGGTGCTCGAGGGCTTTCGCCTCGGCTTCTTTCGCTTG-3' |
| 8 | Helix G-F | 5'-GCGAAAATCTTTATTTTCAAGGGGGGTCTGCAGAGGCGGCTTGGAAGGAATGGGAAGCCAAGGAC-3' |
| 9 | Helix G-R | 5'-GGTGGTGGTGGTGGTGCTCGAGTGCTTTCGCCGCCCAGTCCTTGGCTTCCCATTCC-3' |
| 10 | Helix H-F | 5'-GCGAAAATCTTTATTTTCAAGGGGGGTCTGCCGAACAGGCTTGGAAAGAGTGGGAGGCTAAAGATTG-3' |
| 11 | Helix H-R | 5'-GGTGGTGGTGGTGGTGCTCGAGTGCCTTCACCGCCCAATCTTTAGCCTCCCACTCTTTC-3' |
| 12 | Helix R2-F | 5'-GCGAAAATCTTTATTTTCAAGGGGGGTCTGCAGAGGCGGCTTGGAAGGAAGCAGAAGCCAAGGAC-3' |
| 13 | Helix R2-R | 5'-GGTGGTGGTGGTGGTGCTCGAGTGCTTTCGCCGCCCAGTCCTTGGCTTCTGCTTCC-3' |
| 14 | CPG2-Helix-to-pET15b-F | 5'-GCGGCAGCCATATGGGTTCTTTGGTTGTTGGTGATAATATTGTTGG-3' |
| 15 | CPG2-Helix-to-pET15b-R | 5'-GTTAGCAGCCGGATCCTTATGCTTTATGTTTCCAG-3' |
| 16 | TEV-to-MMP-F | 5'-ACGATCAAAATCAGCAGGTGGTAGCCCTCTTGGTATGTGGTCTCGC-3' |
| 17 | TEV-to-MMP-R | 5'-GCCGCCTCTGCAGACCCGCGAGACCACATACCAAGAG-3' |
| 18 | pQE80L-Linearization-F | 5'-TAAAAGCTTAATTAGCTGAGCTTGGACTCCTG-3' |
| 19 | Domain-Extension-F | 5'-GCAGGTGGTAGCGAAAATCTTTATTTTCAAGGGGGGTCT-3' |
| 20 | Domain-Extension-R | 5'-CTGGATCTATCAACAGGAGTCCAAGCTCAGCTAATTAAGCTTTTA-3' |
| 21 | CPG2$_{\text{CP-N89}}$-TEV-CHis-Linearization-F | 5'-TAAGGATCCGGCTGCTAACAAAGCCC-3' |
| 22 | CPG2$_{\text{CP-N89}}$-TEV-CHis-Linearization-R | 5'-AGACCCCCCTTGAAAATAAAGATTTTCGC-3' |
| 23 | Pro-CPG2-1-Disulfide-1-A177C[†] | 5'-CAGATGAAGAATGCGGTTCATTTGGTTCACGTGATTTGATCC-3' |
| 24 | Pro-CPG2-1-Disulfide-1-M425C[†] | 5'-CCGCAAGATTGAACTGTGCCTGGCGGAAGCGAATC-3' |
| 25 | Pro-CPG2-2-Disulfide-2-L118C[†] | 5'-GGATACAGTTTATTGCAAAGGTATCCTAGCAAAAGCACC-3' |

| Primer Number | Primer Name | Primer Sequence |
|---|---|---|
| 26 | Pro-CPG2-2-Disulfide-2-I432C† | 5'-CTTAAAGCCGCACAGGCCTGCGCGAAAAAATATGGTCTC-3' |
| 27 | Pro-CPG2-2-Disulfide-3-D387C† | 5'-GGGTCTACCTGGTTTTGGTTATCATAGTTGCAAAGCTGAATATG-3' |
| 28 | Pro-CPG2-2-Disulfide-3-A431C† | 5'-CGCTTAAAGCCGCACAGTGCATTGCGAAAAAATATGG-3' |
| 29 | Pro-CPG2-TEV-to_MMP-F | 5'- GTAACACGATCAAAATCAGCAGGTGGTAGCCCTCTTGGTATGTGGTCTCGC-3' |
| 30 | Pro-CPG2-1-TEV-to_MMP-R | 5'-CACCCGTGACATGGTAGGTAATTTGAGACCCGCGAGACCACATACCAAG-3' |
| 31 | Pro-CPG2-2-TEV-to_MMP-R | 5'-GGTTTCATTGCCAATGTGTACAGTAGAAGACCCGCGAGACCACATACCAAG-3' |
| 32 | Pro-CPG2-3-TEV-to_MMP-R | 5'-CGGGCGATGAGTTCTGCCAAGCGAAGAGACCCGCGAGACCACATACCAAG-3' |
| 33 | Pro-Domain-1-F | 5'-CTGGTGCCGCGCGGCAGCCATATGTGGCAAATTACCTACCATGTCACGGGTG-3' |
| 34 | Pro-Domain-2-F | 5'-CTGGTGCCGCGCGGCAGCCATATGTGGTCTACTGTACACATTGGCAATGAAACC-3' |
| 35 | Pro-Domain-3-F | 5'-CTGGTGCCGCGCGGCAGCCATATGTGGCTTCGCTTGGCAGAACTCATCG-3' |
| 36 | Pro-Domains-R | 5'-CTTCCTTTCGGGCTTTGTTAGCAGCC-3' |

†Only the forward primer is given. The reverse primer is the reverse complement.
